# The flavivirus protein NS4B recruits the *cis*-Golgi protein ACBD3 to modify ER-Golgi trafficking for virion release

**DOI:** 10.1101/2024.04.03.587877

**Authors:** Wai-Lok Yau, Marie B. A. Peters, Marie Sorin, Sebastian Rönfeldt, Lauri I. A. Pulkkinen, Lars-Anders Carlson, Anna K. Överby, Richard Lundmark

## Abstract

Flavivirus infection involves extensive remodeling of the endoplasmic reticulum (ER), which is key to both the replication of the viral RNA genome as well as the assembly and release of new virions. Yet, little is known about how viral proteins and host factors cooperatively facilitate such a vast transformation of the ER, and how this influences the different steps of the viral life cycle. In this study, we screened for host proteins that interact with the tick-borne encephalitis virus (TBEV) protein NS4B and found that the top candidates were coupled to trafficking between ER exit sites (ERES) and the Golgi. We characterized the role of ACBD3, one of the identified proteins, in flavivirus infection and show that it interacts with NS4B to promote infection across multiple flavivirus species. Using ACBD3 knockout cells, we found that the depletion of ACBD3 inhibited TBEV replication by preventing the trafficking of virions from the cell. We found that ACBD3 promotes flavivirus infection via a different mechanism than its previously described role in picornavirus infection. ACBD3 was enriched at modified ERES-Golgi contact sites to support virus replication. Therefore, we propose that ACBD3 promotes flavivirus replication by modifying the trafficking between the ERES and the Golgi to enable the release of new virions.

**Author summary:** Flaviviruses including dengue virus and tick-borne encephalitis virus are group of viruses that widely affecting the health of human. During infection, flavivirus particles enter host cells and transform the endoplasmic reticulum (ER), which is the main structure for protein synthesis in cells. New flaviviral particles are produced in the transformed ER and then released to the Golgi apparatus, which is the main structure for protein transport in cells. It is unclear how the particles are transported from the ER to Golgi. Here we screened the factors that interact with viral proteins and identified a Golgi protein called ACBD3 as an important factor supporting flavivirus particles release from ER to Golgi. We showed that ACBD3 is recruited by flavivirus to a modified connection between ER and Golgi for viral particles release from the ER. Our work provides new insights into the fine coordination of virus replication and viral particles transport between organelles inside host cells.

## Introduction

Flaviviruses (genus *Orthoflavivirus*) are enveloped positive-sense single-stranded RNA viruses [1]. This family includes medically relevant members such as dengue viruses (DENV), Zika virus (ZIKV), Japanese encephalitis virus (JEV), yellow fever virus (YFV), West Nile virus (WNV), and tick-borne encephalitis virus (TBEV) [2]. Flavivirus virions enter the host cell via endocytosis which is followed by the fusion of the viral envelope with the endosomal membrane. This releases the genome which is directly translated as single polyprotein at the endoplasmic reticulum (ER). The polyprotein is post-translational processed by host and viral proteases into three structural proteins capsid (C), pre-membrane (prM), and envelope (E), as well as seven non-structural (NS) proteins (NS1, NS2A, NS2B, NS3, NS4A, NS4B, and NS5) [3]. The expression of the viral proteins allows the virus to hijack the cell, and to repurpose the ER and the secretory pathway for the viral genome replication, virion assembly, and export [3–5]. The generation of an extensively transformed ER membrane (TERM) is a hallmark of flavivirus infection and includes small ER membrane invaginations, replication organelles (ROs), where the new genome copies are synthesized [6–13]. The newly synthesized RNA genomes exit the ROs and are either translated to yield more viral proteins or packaged into virions. The genomes that end up packaged interact with the C protein on the ER membrane to form nucleocapsids (NCs) that then bud into the lumen of the ER and acquire the lipid envelope, and the embedded prM and E proteins [6,7,14,15]. This process yields immature virions that need to undergo protease cleavage and pH-mediated conformational changes to produce infectious virions that are then secreted from the cell [16–20].

While TERM formation, and the remodeling of the secretory pathway are crucial for productive flavivirus infection, the details of the process are poorly understood. In particular, the coordinated export of virions, but not of unassembled viral proteins, from the ER to the Golgi remains enigmatic. In healthy cells, anterograde and retrograde trafficking between ER and Golgi is concentrated to ER exit sites (ERES) where the coat protein complex II (COPII) machinery facilitate ER-Golgi trafficking and COPI vesicles mediate transport from Golgi to the ER [21,22]. These processes are intimately coupled and dependent upon each other. Many proteins are involved in regulating this intricate balance of membrane trafficking. Interestingly, recent data show that ERES-Golgi contacts can be modified to facilitate trafficking of large proteins [23–27]. Modification of these contacts has also been described during bacterial infection [28]. Therefore, it is likely that these sites are also exploited by the flaviviral NS proteins by specifically interacting with the proteins that regulate ERES-Golgi contact sites [12,29–31]. The viral protein NS4B is a small integral membrane protein, that interacts both with other viral proteins, as well as host proteins. NS4B seem to play a critical role in the transformation of the ER, and it is proposed to participate in the generation of ROs [29,31,32]. However, NS4B is detected throughout the entire TERM and its precise contribution to membrane remodeling and network of interacting host proteins has not been thoroughly investigated.

In this study we used the medically relevant TBEV and Langat virus (LGTV, a low-pathogenic TBEV-like virus) [1] to investigate the factors involved in generation of TERM during infection. By using ascorbate peroxidase (APEX)-based NS4B interactome screening, we identified that several ERES proteins, including the *cis*-Golgi protein acyl-coenzyme A binding domain containing 3 (ACBD3), were found in close proximity to NS4B. This suggested that NS4B has additional functions in the TERM outside of the ROs. Characterization of the role of ACBD3 showed that it acts as a proviral host factor for multiple flavivirus species by controlling virion trafficking from the ER to the Golgi. Depletion of ACBD3 reduced bulk viral RNA synthesis and virion release and resulted in altered TERM morphology with excessive accumulation of viral proteins at the TERM. Our results suggest that ACBD3 is exploited by flaviviruses to facilitate virion trafficking from the ER to the Golgi.

## Results

### Screen for proteins in close proximity to NS4B identified potential host factors coupled to ERES

To identify host proteins that are in involved the TERM formation, we characterized the TBEV NS4B interactome in uninfected cells, as well as in cells infected with LGTV. To this end, we generated a Flp-In T-REx HEK293 cell line with inducible expression of APEX-tagged NS4B. First, we confirmed that the recombinantly expressed NS4B localizes in the ER of uninfected cells (S1 Fig A) and across the entire TERM in during infection (Fig 1A). We then supplemented both uninfected and infected cells with biotin-phenol and hydrogen peroxide, which induces the biotinylation of the proteins in the vicinity of APEX tag on the NS4B [33]. These proteins were then purified with neutravidin beads and identified using quantitative liquid chromatography-mass spectrometry (LC-MS) analysis (Fig 1B and S1 Figs B-C). We identified 173 proteins that were common for both the infected and uninfected datasets (S1 Table). Interestingly, the ERES proteins ACBD3, TFG, SEC23IP and KTN1 were majorly enriched in the screen (Figs 1C-D) [34–40]. Additionally, 17 other proteins identified in the screen are also involved in ER-Golgi trafficking according to STRING functional enrichment analysis (S1 Table). Furthermore, the transcription factor SMARCA1 was greatly enriched in infected cells as compared to uninfected, suggesting that its close proximity to NS4B is dependent on other viral factors in addition to NS4B (Fig 1C). Finally, NS3 and NS5 were detected in the assay in infected cells, which likely reflects their ability to form complexes with NS4B in the ROs during RNA replication [41–43] (S1 Table).

**Fig 1.**
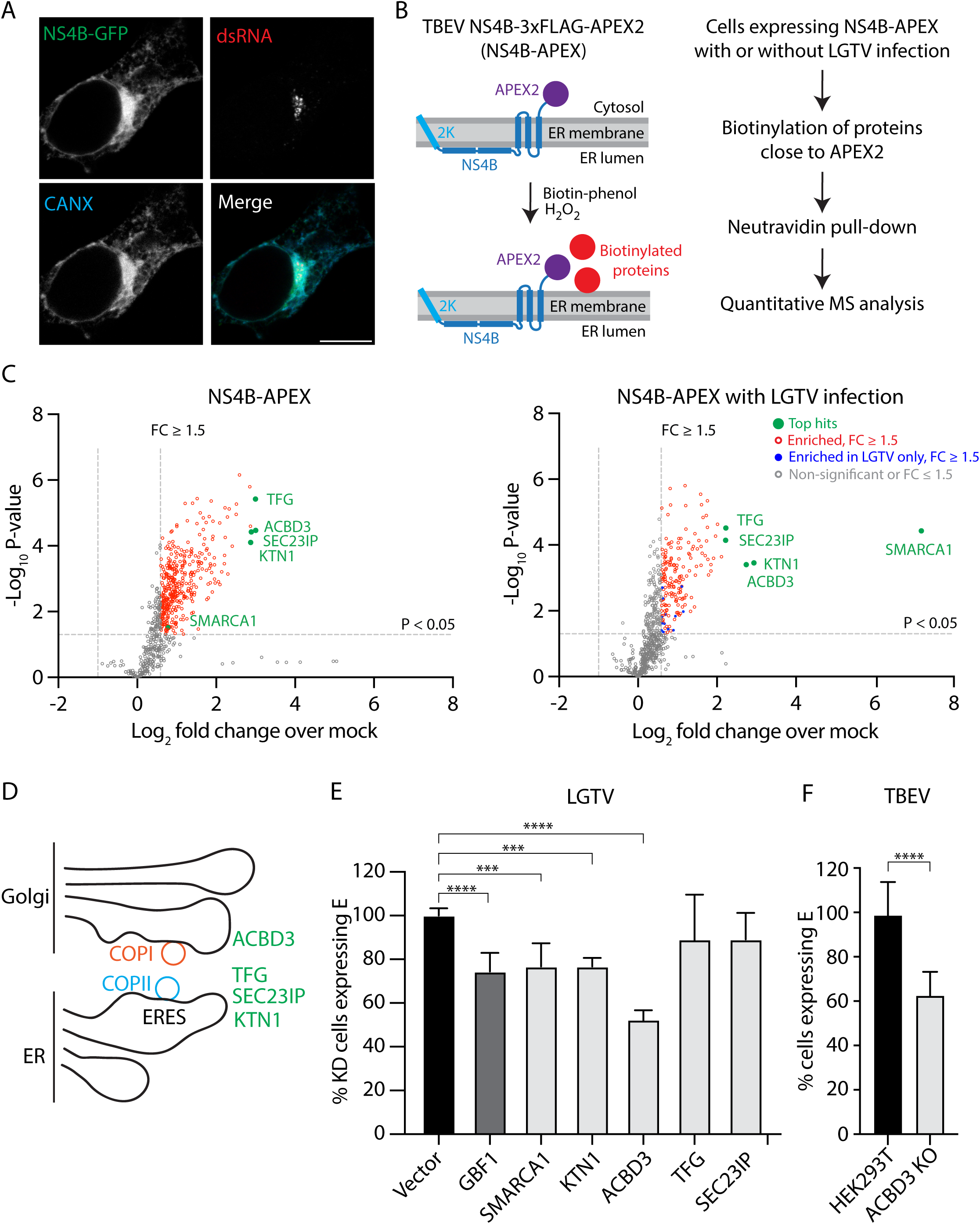
Identification of ACBD3 as a potential host factor in NS4B-APEX proximal protein analysis. (A) Confocal fluorescent micrographs of HEK293T cells transiently expressing NS4B-GFP at 16 h.p.i. with LGTV (MOI 10) and stained with anti-calnexin (CANX) antibodies and anti-dsRNA (dsRNA) antibodies. Scale bar, 10 µm. (B) Schematic illustration of the NS4B-APEX2 proximal protein biotinylation screen used in this study. (C) Volcano plots of the proteins identified in the NS4B-APEX2 proximal protein biotinylation screen. Proteins are indicated by color-coded dots based on the fold change of abundance (FC) values in doxycycline-induced NS4B-APEX cells (left graph) and NS4B-APEX cells infected with LGTV (MOI 10, 16 h.p.i.) (right graph). Statistical significance in p-value was calculated using unpaired t-test. (D) Schematic illustration of ERES-Golgi contacts and the top hit proteins ACBD3, TFG, SEC23IP and KTN1 identified in Fig 1C. (E) Quantification of the percentage of HEK293T cells expressing E protein at 24 h.p.i. with LGTV (MOI 1) after CRISPR-Cas9-mediated knockdown of the indicated proteins. Mean ± SD of 3 independent experiments. One-way ANOVA with Dunnett multiple tests, *** p < 0.005, **** p < 0.0001. (F) Quantification of the percentage of HEK293T cells and ACBD3 KO cells expressing E protein at 24 h.p.i. with TBEV (MOI 0.1). Mean ± SD of at least 10 biological replicates from 2 independent experiments. Unpaired t-tests, **** p < 0.0001.

We moved forward with the top-scoring hits, ACBD3, TFG, SEC23IP, KTN1 and SMARCA1. First, we verified that the proteins co-localized with NS4B-mCherry using confocal microscopy (S1 Figs D-E). To test if the selected hit proteins are important for LGTV infection, we used CRISPR-Cas9 knockdown (KD) to deplete the proteins. The reduction in protein level was confirmed for ACBD3, SMARCA1 and SEC23IP for which antibodies are available (S1 Fig F). As a positive control, cells were depleted of GBF1, which controls Golgi to ER trafficking and was previously shown to inhibit LGTV infection [44]. The infection rates in the KD cell lines were analyzed with flow cytometry by quantifying the fraction of GFP reporter-expressing cells also positive for E after antibody staining. The depletion of GBF1, SMARCA1, KTN1 and ACBD3 significantly reduced the fraction of infected cells compared to vector-only control (Fig 1E). However, the effects of both TFG and SEC23IP KD were not statistically significant (Fig 1E). As ACBD3 KD had the greatest effect on LGTV infection, we followed up on this hit protein using ACBD3 knockout (KO) cells [45] to investigate if it also affected TBEV infection. Indeed, the fraction of cells infected by TBEV, was significantly reduced in cells lacking ACBD3 (Fig 1F).

### The role of ACBD3 in flavivirus infection differs from its proviral function in picornaviruses

As depletion of ACBD3 affected both LGTV and TBEV infection, we used ACBD3 KO cells to investigate if it is needed by all flaviviruses, or only by the tick-borne flavivirus clade [1].

We used a panel of flaviviruses consisting of LGTV, TBEV, ZIKV, JEV, WNV, YFV and DENV serotype 2 (DENV2) and observed that ACBD3 KO reduced the number of infected (dsRNA-positive) cells across the panel, suggesting a genus-level role (Fig 2A).

**Fig 2.**
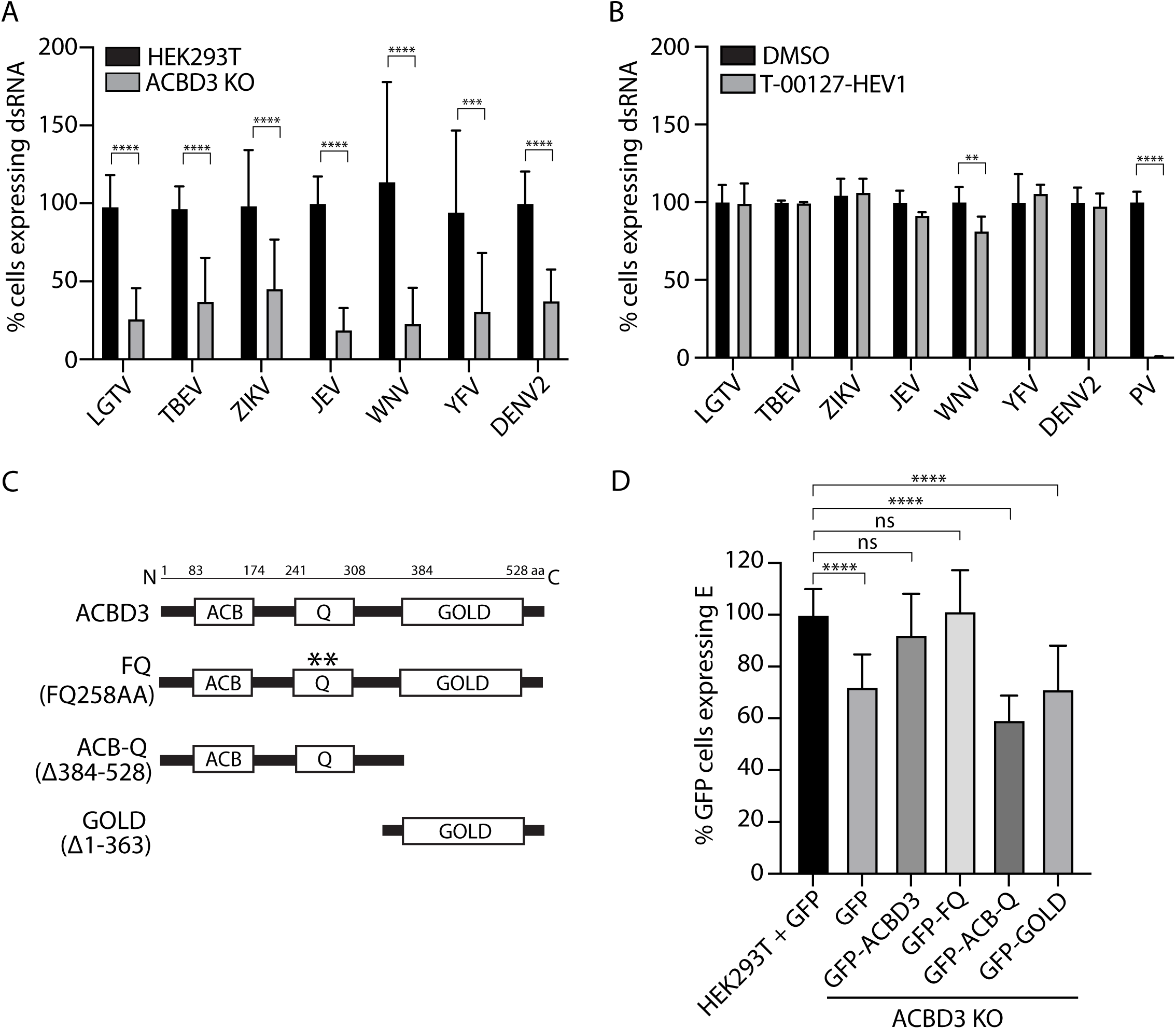
ACBD3 promotes flavivirus infection in a mechanism independent of PI4KB. (A) Quantification of the percentage of HEK293T cells and ACBD3 KO cells positive for dsRNA at 24 h.p.i. with different flaviviruses (MOI 0.1, or 0.5 for DENV2). Data were normalized to the HEK293T control cells. Mean ± SD of at least 6 biological replicates from 2 independent experiments. Unpaired t-test, *** p < 0.001, **** p < 0.0001. (B) Quantification of the percentage of A549 cells positive for dsRNA at 24 h.p.i. with different flaviviruses (MOI 0.1, or 0.5 for DENV2) or at 6 h.p.i. with poliovirus (MOI 10), in the absence and presence of T-00127-HEV1 (1.25 µM) as indicated. Data were normalized to DMSO-treated A549 control cells. Mean ± SD of at least 5 biological replicates from 3 independent experiments. Unpaired t-test, ** p < 0.005, **** p < 0.0001. (C) Schematic illustration of the domains of ACBD3 and the ACBD3 mutants used in the study. (D) Quantification of the percentage of GFP-positive HEK293T cells and ACBD3 KO cells expressing E protein at 24 h.p.i. with TBEV (MOI 0.1) after transient transfection with GFP-tagged protein constructs (24 hours) as indicated. Data were normalized to GFP-positive HEK293T control cells. Mean ± SD of at least 10 biological replicates from 2 independent experiments. One-way ANOVA with Dunnett multiple tests, ns p > 0.05, **** p < 0.0001.

ACBD3 has a known proviral function, as it is an important host factor for picornaviruses. During infection, it recruits PI4KB to RNA replication sites to promote the synthesis of the phosphatidylinositol-4-phosphate (PI4P) lipids that are critical for the function of the picornaviral RNA polymerase [46]. To test if the effects of ACBD3 deletion are the consequence of reduced PI4KB recruitment, we used the PI4KB inhibitor T-00127-HEV1 with a strong antiviral effect on picornaviruses [47]. The treatment had no effect on cell viability up to 5 µM, and as previously reported, it reduced poliovirus (PV) infection in a dose-dependent manner with near-complete inhibition observed at 1.25 µM (S2 Figs A-B). However, at 1.25 µM, the compound did not inhibit LGTV, TBEV, ZIKV, JEV, YFV, or DENV2 replication (Fig 2B). Interestingly, a small but significant decrease in the number of WNV-infected cells was detected. Nonetheless, for the majority of flaviviruses, PI4KB does not seem to be important for infection, and therefore the role of ACBD3 seems to be independent of PI4KB recruitment.

Next, we investigated which ACBD3 domain(s) are needed for the protein’s proviral functions by transfecting ACBD3 KO cells with constructs expressing GFP-tagged wild-type (WT) or mutant ACBD3 and quantifying the fraction of TBEV-infected cells. ACBD3 is composed of an ACB domain (ACB) that binds acyl-CoA, a Q domain that interacts with PI4KB and a Golgi dynamics domain (GOLD) which facilitates various membrane and protein interactions [48]. Transfection with full-length GFP-tagged ACBD3 (WT) rescued the infection, but transfection of either the GOLD domain on its own, or the ACB-Q construct lacking the GOLD domain did not (Figs 2C-D). Interestingly, productive infection was also restored by complementation with the FQ mutant that lacks the PI4KB binding site [46], further confirming that ACBD3 promotes flavivirus and picornavirus infections with different mechanisms (Fig 2D).

### ACBD3 depletion leads to less bulk RNA replication and reduced virion release

To pinpoint which part of the viral life cycle ACBD3 plays a role in, we investigated the differences in viral protein and RNA production, as well as viral titer between WT and KO cells. When infected with LGTV at multiplicity of infection (MOI) 1, the amounts of viral proteins NS3, E, and prM were lower than in control cells (S3 Fig). Indeed, when analyzed by fluorescent light microscopy, the number of E-positive cells was smaller in KO cells (Figs 3A-B). However, we observed that in the KO cells that did get infected, the E protein signal seemed more intense than in WT cells (Fig 3A). When we quantified the viral titers produced in KO and WT cells at different MOIs using focus forming assay, the viral titers were consistently significantly lower in KO cells (Fig 3C). Next, we used quantitative reverse-transcription PCR (RT-qPCR) to assay the effects of ACBD3 KO on bulk viral RNA production. At MOI 1, the bulk viral RNA in KO cells was significantly lower than in WT cells at 8, 12, and 16 hours post infection (h.p.i.) (Fig 3D).

**Fig 3.**
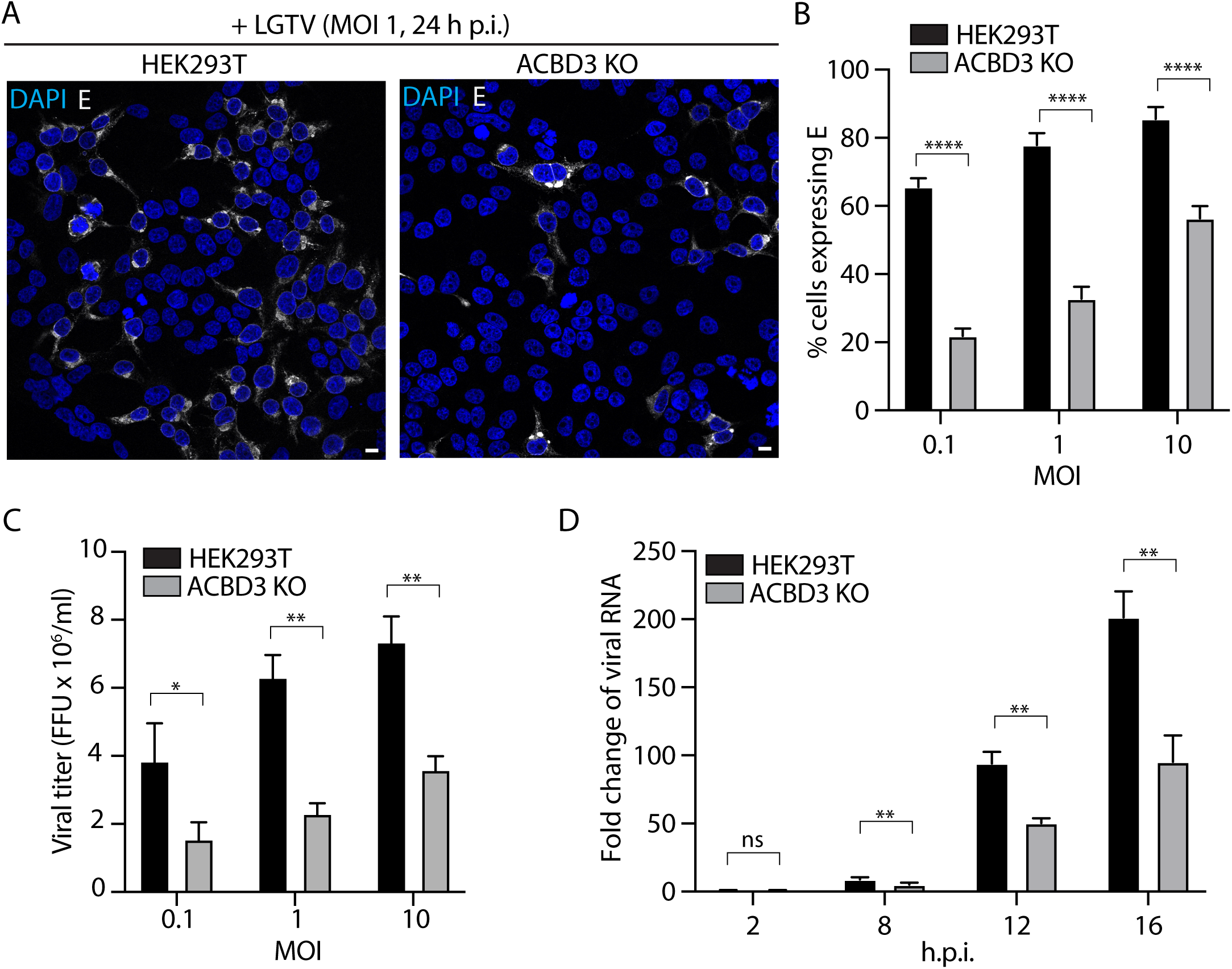
Depletion of ACBD3 inhibits intracellular amplification and release of LGTV. (A) Confocal fluorescent micrographs of LGTV-infected (MOI 1, 24 h.p.i.) HEK293T cells and ACBD3 KO cells stained with anti-E antibodies. Scale bar, 10 µm. (B) Quantification of the percentage of HEK293T cells and ACBD3 KO cells expressing E proteins at 24 h.p.i. with LGTV (MOI 0.1, 1 or 10). Mean ± SD of at least 7 biological replicates. Unpaired t-test, **** p < 0.0001. (C) Quantification of viral titer of the supernatant of LGTV-infected (24 h.p.i.) HEK293T cells and ACBD3 KO cells by focus forming assay. Mean ± SD of 4 biological replicates. Unpaired t-test, * p < 0.05, ** p < 0.005. (D) Quantification of viral RNA of LGTV-infected (MOI 1) of HEK293T cells and ACBD3 KO cells by RT-qPCR. Fold change of actin-normalized viral RNA was calculated by comparing with normalized viral RNA input at 2 h.p.i. Mean ± SD of 3 biological replicates. Unpaired t-test, ** p < 0.005.

### Loss of ACBD3 leads to altered TERM morphology

Next, we focused on the ACBD3 KO cells that did get infected and expressed viral proteins. Fluorescence microscopy analysis revealed that in the ACBD3 KO cells, both E and NS3 were focused as bright foci that were not observed in the WT cells (Fig 4A). To quantify this phenomenon, we determined the regions in cells where the NS3 fluorescence intensity was continuously above a set threshold and the detected area was larger than 10 μm^2^ (Figs 4B-C). Then, we quantified the mean fluorescence intensity of NS3 and E within these NS3-positive areas. Notably, the mean fluorescent intensities of NS3 and E within the NS3 regions were significantly higher for the KO than the WT cells (Figs 4D-E). Similar analysis of dsRNA showed that the mean intensity of dsRNA was also higher in the NS3 regions of ACBD3 KO cells than in WT cells (Fig 4F). This showed that the replication of viral RNA and viral protein translation are still possible in ACBD3 KO cells, but that the viral components are abnormally accumulating due to the effects of ACBD3 KO on the TERM.

**Fig 4.**
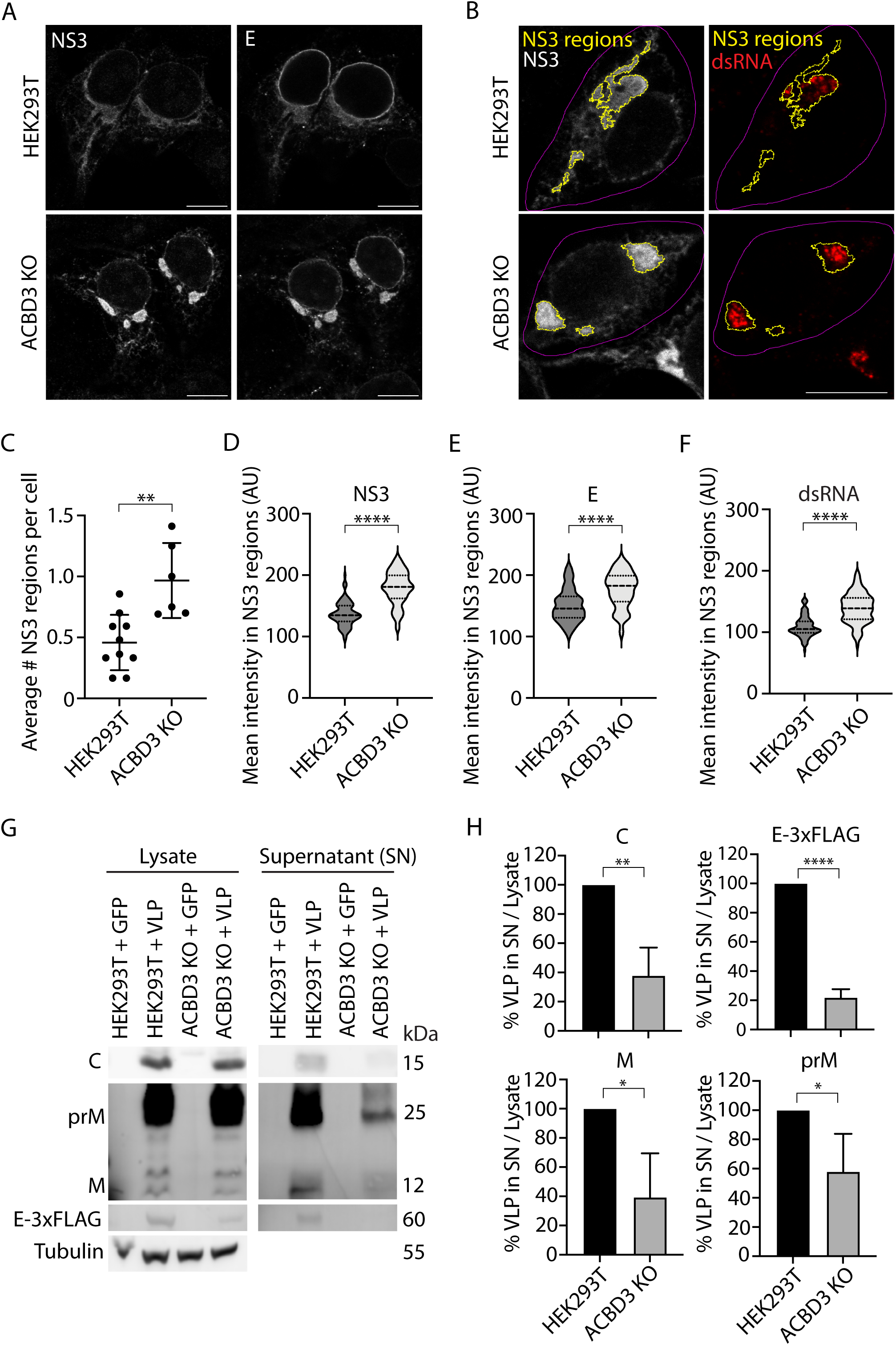
Depletion of ACBD3 leads to the accumulation of NS3, E and dsRNA. (A) Confocal fluorescent micrographs of LGTV-infected (MOI 1, 16 h.p.i.) HEK293T cells and ACBD3 KO cells stained with anti-NS3 antibodies and anti-E antibodies. Scale bar, 10 µm. (B) Confocal fluorescent micrographs of LGTV-infected (MOI 1, 16 h.p.i.) HEK293T cells and ACBD3 KO cells stained with anti-NS3 antibodies and anti-dsRNA antibodies. Yellow lines represent the identified NS3 regions (size > 10 µm^2^) with a continuous fluorescence intensity of NS3 over the set threshold (see Materials and methods). Magenta lines represent the manually assigned outline of NS3-positive cells. Scale bar, 10 µm. (C) Quantification of the average number of NS3 regions (> 10 µm^2^) per cell. Data shown as mean ± SD. Number of cells analyzed, HEK293T (n = 215 in 10 imaged areas), ACBD3 KO (n = 108 in 6 imaged areas). Each data point represents one imaged area of the slides. Unpaired t-test, ** p < 0.005. (D and E) Quantification of the mean fluorescence intensities of NS3 (D) and E (E) within the identified NS3 regions. Data are presented in violin plots with median and quartiles indicated by dashed and dotted lines, respectively. Number of NS3 regions analyzed, HEK293T (n = 95), ACBD3 KO (n = 103). Unpaired t-test, **** p < 0.0001. (F) Quantification of mean fluorescence intensity of dsRNA within the identified NS3 regions. Data are presented in violin plot with median and quartiles. Number of NS3 regions analyzed, HEK293T (n = 99), ACBD3 KO (n = 104). Unpaired t-test, **** p < 0.0001. (G and H) VLP secretion of HEK293T cells and ACBD3 KO cells transiently co-transfected with C, prM and E-3xFLAG (24 hours) analyzed with western blot using specific antibodies to C, prM and 3xFLAG and quantified in Fig 4H. Data were normalized with HEK293T + VLP control. Mean ± SD of 3 biological replicates. Unpaired t-test, * p < 0.05, ** p < 0.005, **** p < 0.0001.

### Virus-like particle secretion is impaired in ACBD3 KO cells

ACBD3 is located in the *cis*-Golgi and plays a key role in protein trafficking by modifying the connection sites between ERES and the *cis*-Golgi [39,40]. This suggested that modulation of trafficking might be related to the role of ACBD3 in flavivirus infection. To investigate this, we used a virus-like particle (VLP) secretion assay to decouple the possible effects of ACBD3 KO on genome replication from those affecting virion assembly and egress. We transfected ACBD3 KO cells with plasmids expressing the viral C, prM, and E-3xFLAG proteins, which results in the secretion of VLPs consisting of these proteins [44]. Immunoblot analysis showed that while WT and ACBD3 KO cells produced similar amounts of viral proteins, the amount of viral proteins (in the form of VLPs) secreted into the supernatant was significantly lower in KO than WT cells (Figs 4G-H). This showed that the loss of ACBD3 results in either a defective VLP assembly or protein trafficking.

### ACBD3 is located at contact sites between TERM and Golgi

To characterize the ACBD3 localization in detail, we used confocal microscopy which showed that ACBD3 almost completely co-localized with the *cis*-Golgi marker GM130 in both control and infected cells (S5 Figs A and C). Furthermore, SEC23IP, another ERES hit in the screen, was detected in discrete puncta overlapping with GM130 in agreement with that connections might form between ERES and *cis*-Golgi. This was observed in both infected and uninfected cells (S5 Figs B and D), suggesting that while NS4B may localize to these sites during infection, it is not responsible for generating them. Additionally, ACBD3 co-localized with both the NS3 and E, as the signal for both viral proteins was detected in a large area (S5 Figs C and E). However, ACBD3 was not found at the sites of dsRNA replication. Instead, ACBD3 was located next to these locations (S5 Fig C), suggesting that ACBD3 is not directly involved in RO formation or RNA replication, but rather in the coupling of the *cis*-Golgi near these sites.

To resolve the organization of the ERES-Golgi sites, we used structured illumination microscopy (SIM). Interestingly, analysis of the localization of GFP-ACBD3 and SEC23IP, a component of the COPII protein coat, showed that that the punctate ERES positive for SEC23IP overlapped with interconnecting membrane tubules positive for GFP-ACBD3 (Fig 5A and S1 movies A-C). This showed that ACBD3 indeed connects the *cis*-Golgi to the ERES. Similar analysis of infected cells showed that the TERM positive for E protein was composed of a tubular network. SEC23IP was detected as distinct puncta on TERM tubules, and they were connected to ACBD3-positive membrane tubules (Fig 5B and S2 movies A-C). Taken together these data suggested that ACBD3-positive ERES-Golgi contacts are controlling the export of virions.

**Fig 5.**
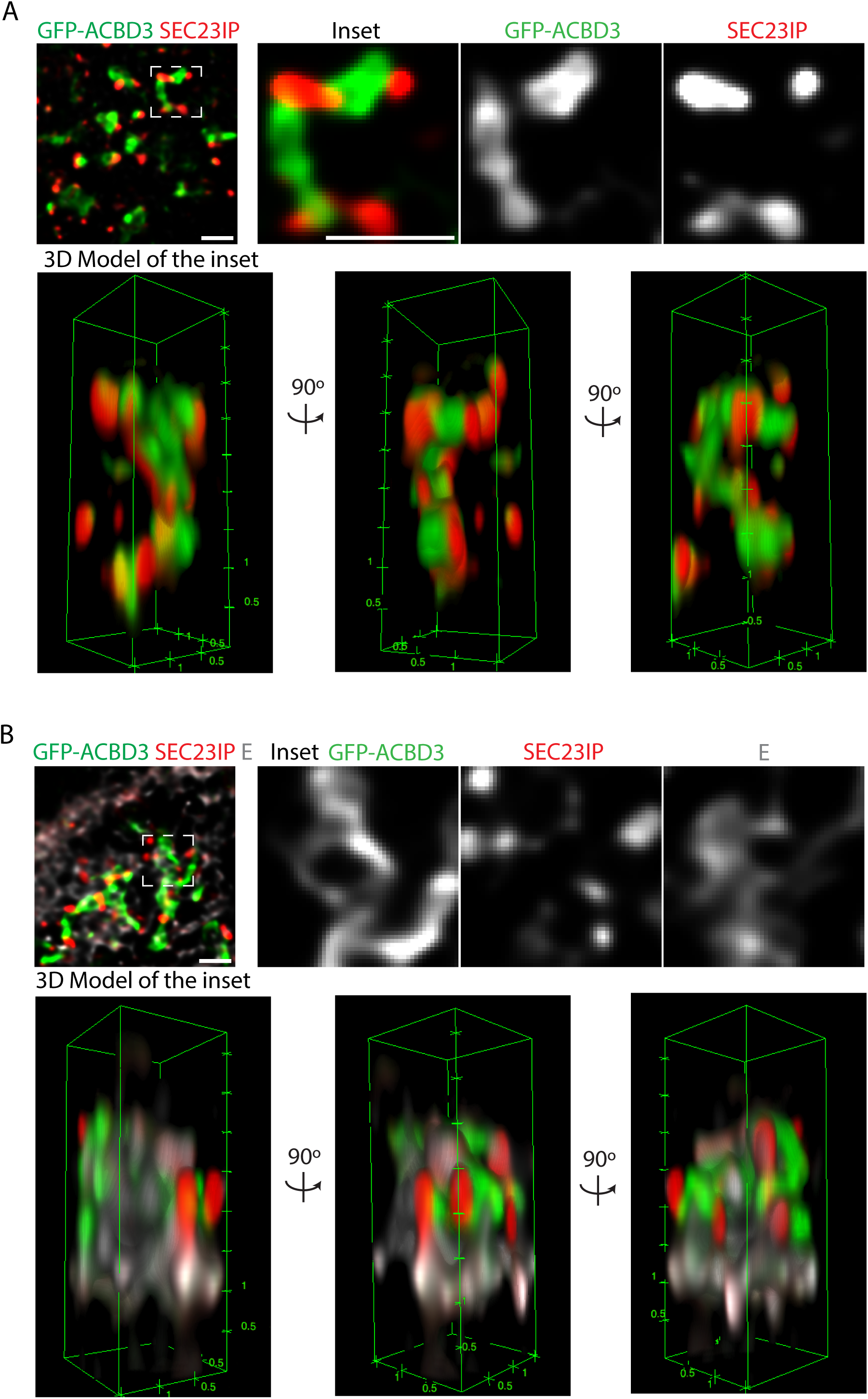
Super-resolution analysis showing close localization of ACBD3 and SEC23IP at the ERES-Golgi contact. (A) SIM fluorescence micrographs at 100 nm resolution of HEK293T cells transiently expressing GFP-ACBD3 and stained with anti-SEC23IP antibodies. Scale bars, 1 µm. Bottom panels show the 3D-representations of the volume SIM imaging data of the inset at 0°, 90° and 180° rotations. (B) SIM fluorescent micrographs at 100 nm resolution of HEK293T cells transiently expressing GFP-ACBD3 at 16 h.p.i. with LGTV (MOI 1) and stained with anti-SEC23IP antibodies and anti-E antibodies. Scale bars, 1 µm. Bottom panels show the 3D-representations of the volume SIM imaging data of the inset at 0°, 90° and 180° rotations.

### ACBD3 is recruited to the TERM via the GOLD domain

To address which domain in ACBD3 that was targeted by NS4B in order to modify the contacts between ERES and Golgi, we used confocal microscopy. Expression of GFP-tagged truncated protein constructs in ACBD3 KO cells showed that the GFP-GOLD domain co-localized with the *cis*-Golgi marker GM130 similar to GFP-ACBD3 (Fig 6A). However, GFP-ACB-Q was primarily detected in the cytosol showing that the GOLD mediates membrane targeting (S6 Fig). To visualize the localization of ACBD3 in relation to the TERM, the co-localization between NS3 and GFP-ACBD3 and GFP-GOLD was analyzed in LGTV-infected cells. In contrast to GFP-ACBD3, the GFP-GOLD was not only detected at the Golgi associated with the TERM, but it localized to the entire NS3-positive area (Fig 6A). Quantification of the NS3 area that was positive for either GFP-GOLD or GFP-ACBD3 showed that the area was on average three times larger for GOLD (Fig 6B). This suggested that the GOLD was directly recruited by NS4B during infection, but that without the other domains the specific connection to ERES-Golgi sites was lost. To test if NS4B was sufficient to recruit the GOLD, we co-expressed GFP-GOLD and NS4B-mCherry in ACBD3 KO cells. Fluorescence microscopy analysis showed that the majority of the NS4B-mCherry co-localized with GFP-GOLD supporting that NS4B can recruit the GOLD to the ER (Figs 6C-D). Taken together, our data support that the expression of NS4B during TBEV infection recruits ACBD3 via the GOLD to modify ERES-Golgi contacts which support efficient release of virions.

**Fig 6.**
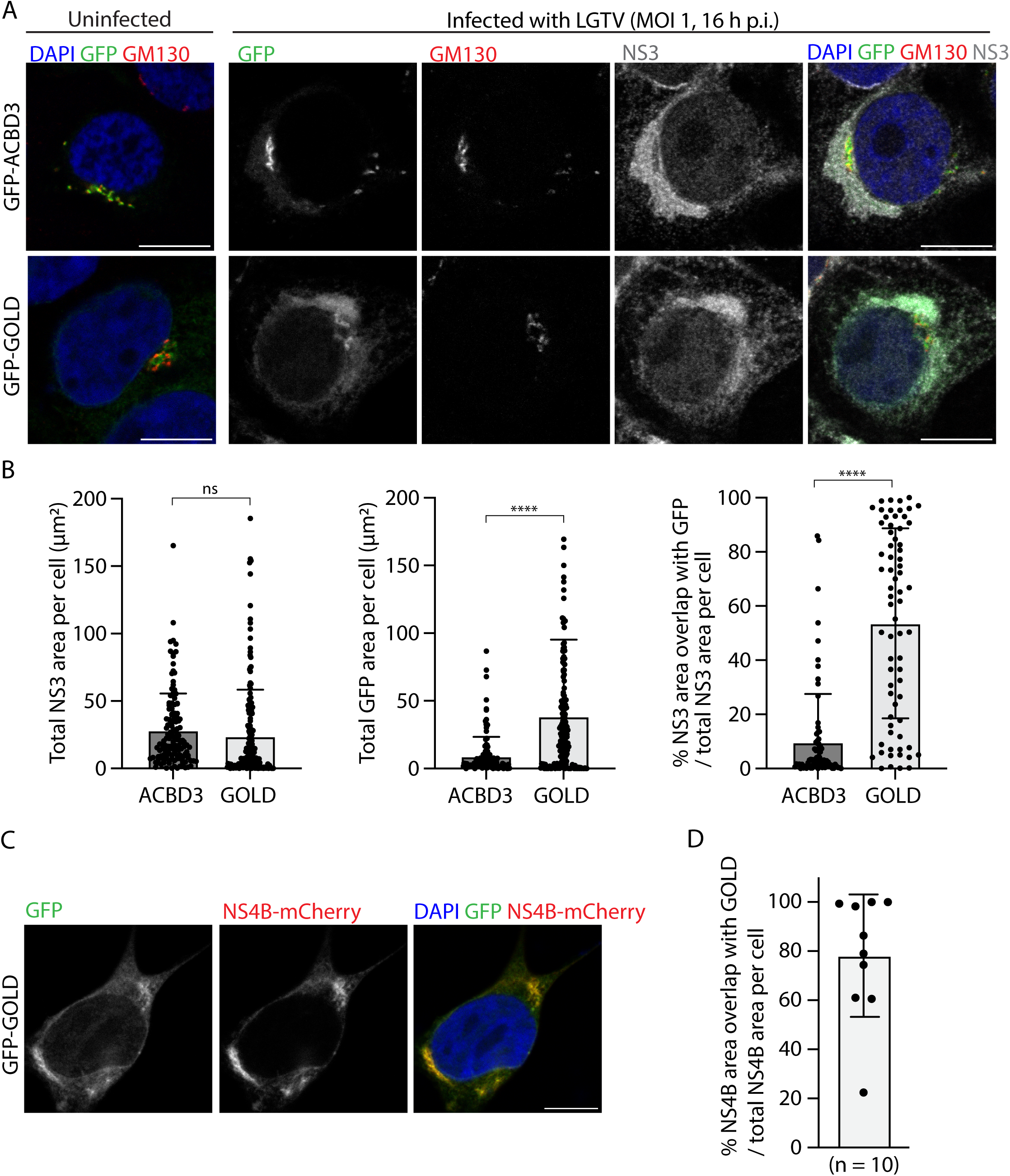
ACBD3 GOLD domain accumulates in the ER in LGTV-infected cells. (A) Confocal fluorescence micrographs of ACBD3 KO cells transiently over-expressing GFP-ACBD3 or GFP-GOLD at 16 h.p.i. with LGTV (MOI 1) and stained with anti-GM130 antibodies and anti-NS3 antibodies. Scale bar, 10 µm. (B) Quantification of the percentage of the fluorescence NS3 area overlapping with GFP-ACBD3 or GFP-GOLD. Data were presented in scatter plots with mean ± SD. Number of NS3 area analyzed, GFP-ACBD3 (n = 130), GFP-GOLD (n = 179). Mann-Whitney test, **** p < 0.0001, ns p > 0.05. (C) Confocal fluorescence micrographs of ACBD3 KO cells transiently expressing NS4B-mCherry and GFP-ACBD3 or GFP-GOLD. Scale bar, 10 µm. (D) Quantification of the percentage of the fluorescence NS4B area overlapping with the fluorescence of GFP-GOLD. Data are presented in a scatter plot with mean ± SD. Number of cells analyzed, n = 10.

## Discussion

The general characteristics of the flavivirus maturation and egress pathway were elucidated decades ago by groundbreaking biochemical and structural work on TBEV and DENV [30,49–51]. However, the model is built on the assumption that the ER and the Golgi function similarly in infected and healthy cells. Instead, electron and light microscopy observations from multiple flavivirus species show that both the ER and the Golgi, as well as the cellular lipidome are extensively remodeled during infection [6,7,11–13,52–55]. Therefore, it is likely that the maturation and secretion of flaviviral particles is more complex than previously thought, especially since recent work has demonstrated that also in uninfected cells, ER-Golgi trafficking includes previously undescribed mechanisms [56–59]. The established model for maturation of virions has also been challenged by the recent high-resolution cryogenic electron microscopy reconstructions of immature flaviviruses. These structures show that the previous model is too simplistic, as steric clashes would occur if the maturation were to proceed as previously thought [16–18,20].

Our APEX screen of host factors that interact with NS4B during flavivirus infection identified 173 candidate proteins with 20 of the hits implicated with ER to Golgi transport. Furthermore, four of the five top hits (ACBD3, TFG, SEC23IP, and KTN1) are associated with ERES [34–40]. This suggested that NS4B might play additional roles outside of the ROs. We confirmed the co-localization of the five top hits with NS4B using fluorescence microscopy and followed up on ACBD3 since it had the greatest effect on LGTV infection in our screen. Our results show that ACBD3 promotes productive infection across multiple flavivirus species, and that the full-length protein (opposed to single domains) is needed for this function.

ACBD3 is a well-characterized proviral factor for picornaviruses, which are non-enveloped viruses that replicate on cytoplasmic membrane vesicles derived from the ER. Therefore, we investigated if ACBD3 performs a similar function during flavivirus infection. The picornavirus protein 3A coordinates ACBD3 to the sites of RNA replication, so it can further recruit PI4KB [46]. The viral RNA polymerase in turn requires the PI4P lipids synthesized by PI4KB for efficient function [60]. The ACBD3 Q domain is responsible for the PI4KB recruitment [46]. In this study we showed that complementation with full-length ACBD3 containing a FQ mutation, which resulted in defective PI4KB binding [46], did revert normal flavivirus infection. This suggests that ACBD3 has a different mechanism in flaviviruses compared to picornavirus infection. We confirmed that PI4KB, and therefore its recruitment by ACBD3 is not critical for flaviviruses by using T-00127-HEV1, a compound that inhibits PI4KB function and therefore prevents picornavirus infection [47]. As previously shown, T-00127-HEV1 treatment inhibited PV replication. However, it did not have an effect across a panel of flaviviruses, apart from a minor reduction in the number of WNV-infected cells. Therefore, we concluded that the main ACBD3 function in flavivirus infection is not related to PI4KB recruitment or function. This is not surprising considering that unlike in the ER-centric flavivirus replication cycle, picornaviral RNA synthesis and assembly occur on cytoplasmic membrane vesicles and virion release is mediated via autophagosomes [61–63]. However, it is intriguing that ACBD3 is hijacked by both distantly related viral taxa. Maybe modulation of or co-opting ACBD3 activity is a prerequisite for any virus that needs to distort the intracellular membranes for its replication.

As ACBD3 functions in ER-Golgi trafficking, we investigated the effects of ACBD3 KO on viral egress. The VLP secretion assay directly showed that ACBD3 knockout reduces the amount of exported viral proteins, which suggests that ACBD3 KO affects either the assembly or secretion of VLPs (and therefore virions). While it is difficult to pinpoint if the KO of ACBD3 dysregulates all export, or purely specific cargoes, it is clear that the pathways relevant for flavivirus virion egress are affected. The other observations are also consistent with trafficking defects, as in the infected ACBD3 KO cells, we observed higher E protein fluorescence intensity than in WT cells. This shows the abnormal accumulation of viral structural proteins in the cell. Furthermore, while we observed lower bulk RNA production in KO cell cultures compared to WT, the infected cells had higher dsRNA fluorescence intensity in KO cells showing that RNA synthesis was not impaired on the cellular level. Instead, both the lower total RNA yield as well as the lower number of E protein-producing cells can be explained by virion assembly or secretion defects that limit subsequent infection rounds in MOI 1 infections. Additionally, the lower infection rates and bulk RNA production may indicate ER or Golgi stress caused by the accumulation of potentially misfolded viral proteins resulting in the clearing of the viral infection in a larger fraction of KO than WT cells as both ER and Golgi stress have been reported to have unclear roles in flavivirus infection [64–66]. Finally, we cannot rule out that ACBD3 KO affects the secretion of NS1, or the interferon response, both of which may modulate how susceptible the cell culture to infection [39,45,67,68].

As our top hits, alongside with 20 others relate to ER-Golgi trafficking, it seems that NS4B is responsible for recruiting host factors to control ER-Golgi trafficking. Traditionally this has been thought to be mediated by COPII vesicular trafficking [21,37,69]. Yet, non-canonical mechanisms for transport between these compartments have recently been described. Analysis of trafficking dynamics in live cells demonstrated that bidirectional transport portals are formed between the ERES and arrival sites at the Golgi [58]. Further, ultrastructural analysis revealed a tubular membrane network extending from the COPII-positive ER to the COPI-positive *cis*-Golgi [59]. Interestingly, ACBD3 has also been involved in modifying ERES-Golgi contacts to mediate Golgi transport of the STING receptor during bacterial infection [39,70]. Based on our studies of ACBD3, we propose a model where NS4B-mediated recruitment of ACBD3, SEC23IP, KTN1 and TFG modifies ERES-Golgi contacts to facilitate efficient trafficking of immature virions from the TERM to the Golgi (Fig 7). This would enable controlled transport of the relatively large virions, while at the same time ensuring that viral proteins remain concentrated in the TERM. Whether the modified trafficking process involves fusion of ER and Golgi membranes into tubular portals or whether remains to be elucidated. Also, the exact mechanism by which the ERES-Golgi contacts are modified by the identified host proteins is not yet clear. In the conventional secretory pathway, ACBD3 has been proposed to control the retrograde trafficking of KDEL receptors (KDELR) from the Golgi to ER. ACBD3 forms a complex with KDELR and depletion of ACBD3 resulted in an accumulation of KDELR at ER and disruption of KDELR cycling [71,72]. Interestingly, both prM and E on the immature virion surface of DENV and JEV has been shown to bind to KDELR [73,74]. It remains to be elucidated if KDELR is directly involved in the ACBD3-mediated transport of TBEV. In summary, this study identified ACBD3 as a novel host factor in flavivirus infection. We propose that ACBD3 functions at ERES-Golgi contact and exploited by flavivirus to mediate trafficking from the TERM to the Golgi.

**Fig 7.**
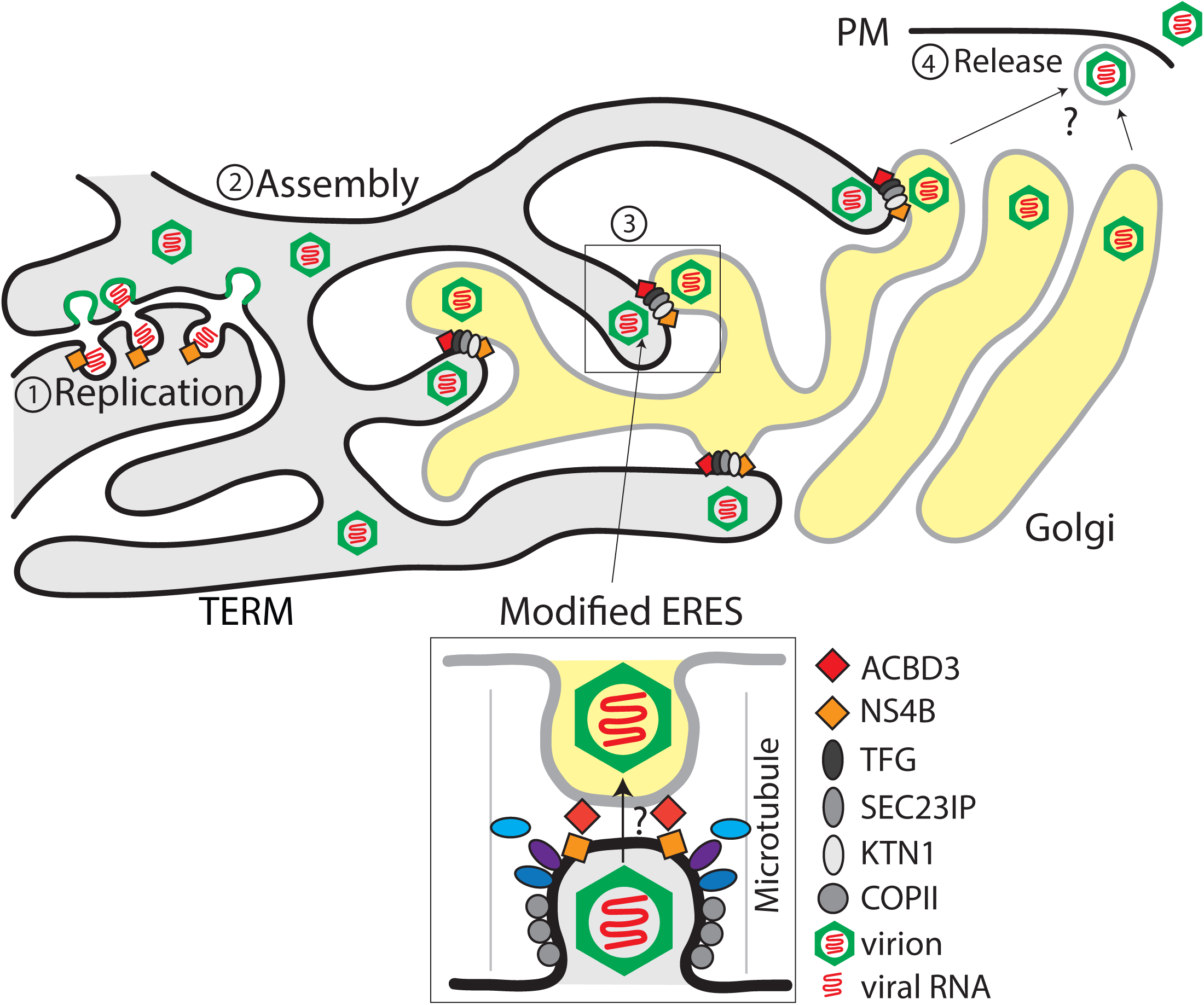
Model of TERM and ACBD3 function in flavivirus infection. Schematic illustration of the proposed model for how NS4B-mediated modification of ERES by recruitment of ACBD3, SEC23IP, TFG and KTN1 directly link the TERM to the *cis*-Golgi in order to promote secretion of virions. Viral ssRNA genomes are replicated in TERM-derived ROs which contain NS4B and other viral and host proteins (1), and the genomes interact with viral structural proteins to form an immature virion that buds into the lumen of the TERM (2). The virions are trafficked to the modified ERES-Golgi contact sites that have been hijacked by NS4B interracting with host proteins to facilitate the transport of large viral cargoes to the Golgi (3). In the Golgi, it is unknown if the virions are trafficked through the Golgi like canonical exocytic cargoes, or if they are transported straight to the PM for release (4).

## Materials and methods

### Cell lines and virus strains

HEK293T cells, HEK293T ACBD3 KO cells and A549 cells were cultured in Dulbecco’s modified Eagle medium (DMEM) supplemented with 10% FBS (Gibco, Thermo Fisher Scientific) at 37°C, 5% CO2. HEK293T ACBD3 KO cells was a gift from Frank van Kuppeveld [45]. Viral stocks were propagated in Vero B4 cells or A549 MAVS knockout cells (Abcam, cat no. ab282354). LGTV strain TP21 (kindly provided by Gerhard Dobler, Bundeswehr Institute of Microbiology, Munich, Germany) was propagated in A549 MAVS knockout cells. TBEV strain Töro-2003 [75] was propagated in Vero B4 cells. Additional virus strains (ZIKV, JEV, DENV, YFV, WNV, PV) used in the study are listed in table 1. Viral titers of flaviviruses were determined by focus forming assay [9] and viral titer of poliovirus was determined by plaque assay (described in the following section).

**Table 1.**
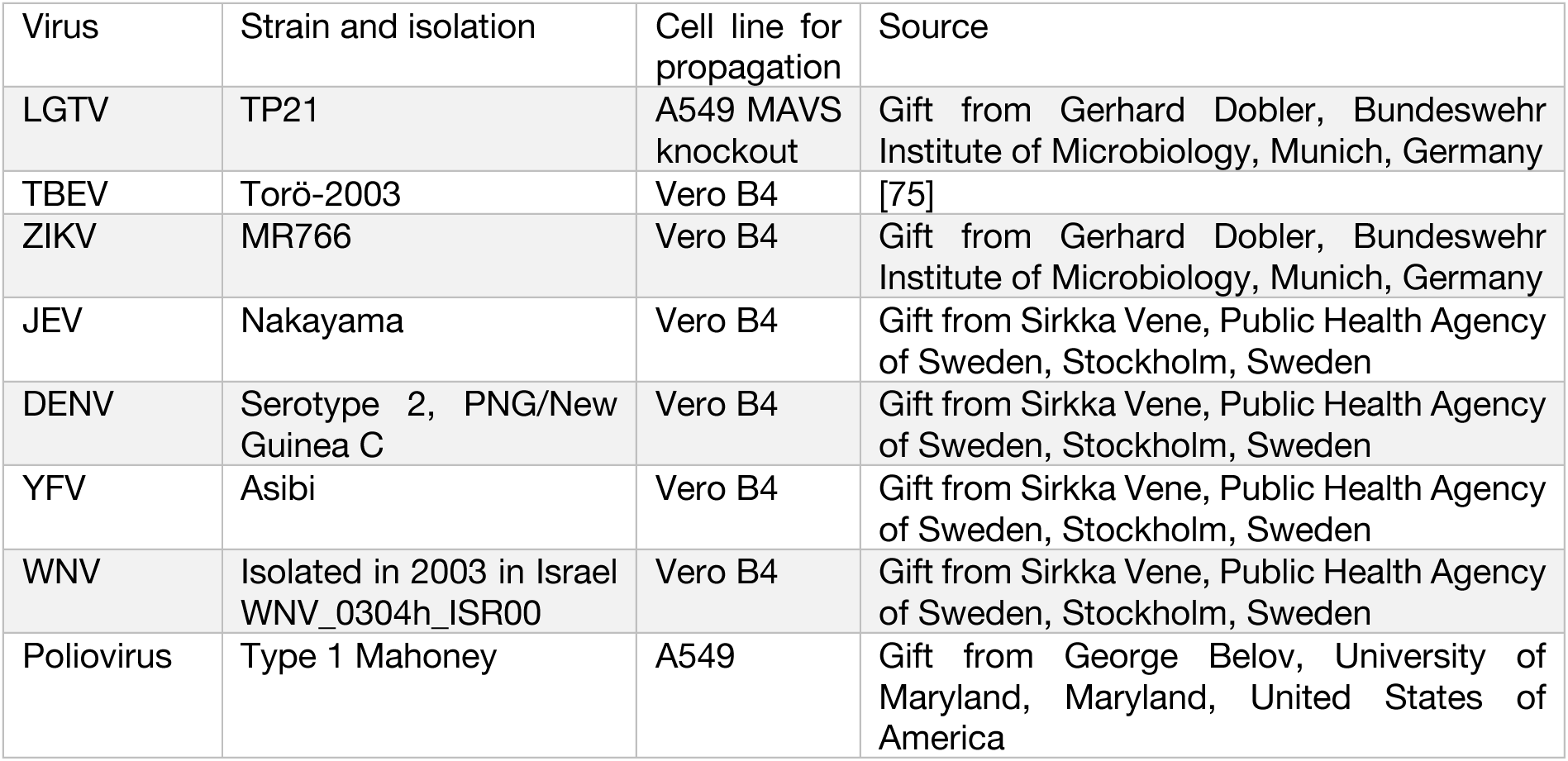
Virus strains used in the study.

| Virus | Strain and isolation | Cell line for propagation | Source |
| --- | --- | --- | --- |
| LGTV | TP21 | A549 MAVS knockout | Gift from Gerhard Dobler, Bundeswehr Institute of Microbiology, Munich, Germany |
| TBEV | Torö-2003 | Vero B4 | [75] |
| ZIKV | MR766 | Vero B4 | Gift from Gerhard Dobler, Bundeswehr Institute of Microbiology, Munich, Germany |
| JEV | Nakayama | Vero B4 | Gift from Sirkka Vene, Public Health Agency of Sweden, Stockholm, Sweden |
| DENV | Serotype 2, PNG/New Guinea C | Vero B4 | Gift from Sirkka Vene, Public Health Agency of Sweden, Stockholm, Sweden |
| YFV | Asibi | Vero B4 | Gift from Sirkka Vene, Public Health Agency of Sweden, Stockholm, Sweden |
| WNV | Isolated in 2003 in Israel WNV_0304h_ISR00 | Vero B4 | Gift from Sirkka Vene, Public Health Agency of Sweden, Stockholm, Sweden |
| Poliovirus | Type 1 Mahoney | A549 | Gift from George Belov, University of Maryland, Maryland, United States of America |

### Focus forming assay

Vero B4 cells in 96-well plates were infected with serial diluted virus samples (supernatant of infected culture) in DMEM for 2 hours at 37°C, 5% CO2. The inocula were then replaced with DMEM supplemented with 2% FBS and 1.2% Avicel (FMC BioPolymer). After 48 h.p.i., the cells were fixed in 4% formaldehyde for 20 min, and then permeabilized with 0.5% Triton X-100 in PBS. The cells were then incubated with anti-E antibodies for 1 hour at RT. The cells were then incubated with antibodies HRP conjugated secondary antibodies for 1 hour at RT. Chromogenic peroxidase substrate Trueblue (KPL, Seracare) was added to the plates for focus visualization. The plates were imaged with a Cytation 5 plate imager (Biotek) and the foci were counted manually. The viral titer was expressed as focus forming units per ml (FFU/ml).

### Plaque assay

Confluent HeLa cells in 6-well plates were infected with serial diluted virus samples (supernatant of infected culture) in minimum essential medium (MEM) for 1 hour at 37°C, 5% CO2. The inocula were then replaced with MEM supplemented with 2% FBS and 0.3% agarose. After 48 h.p.i., the cells were fixed with 5% formaldehyde for 24 hours. The cells were then rinsed with water and stained with 3% crystal violet for plaque visualization. The stained cells were then washed with water. The plaques were counted manually. The viral titer was expressed as plaque forming units per ml (PFU/ml).

### Antibodies, probes, and plasmids

The antibodies and probes used in the study are listed in table 2. The plasmids used in the study are listed in table 3.

**Table 2.** Antibodies and probes used in the study.

| Antibody/Probe | Dilution | Source | Identifier |
| --- | --- | --- | --- |
| Anti-CANX | 1:50000 (WB),<br>1:1000 (IF) | Abcam | Cat# ab22595 |
| Anti-ACBD3 | 1:2000 (WB),<br>1:1000 (IF) | Atlas Antibodies | Cat# HPA015594 |
| Anti-GM130 | 1:300 (IF) | BD Transduction | Cat# 610823 |
| Anti-SMARCA1 | 1:500 (IF) | Atlas Antibodies | Cat# HPA064712 |
| Anti-SEC23IP | 1:500 (IF) | Atlas Antibodies | Cat# HPA043305 |
| Anti-GAPDH | 1:5000 (WB) | Merck Millipore | Cat# MAB374 |
| Anti-Tubulin | 1:4000 (WB) | Abcam | Cat# ab6046 |
| Anti-dsRNA clone J2 | 1:1000 (IF) | Scicons, Nordic MUBio | Cat# 10010500 |
| Anti-NS3 | 1:1000 (IF),<br>1:5000 (WB) | [76] |  |
| Anti-C | 1:1000 (WB) | [44] |  |
| Anti-M | 1:1000 (WB) | [77] |  |
| Anti-E clone 1786.3 | 1:1000 (IF) | [78] |  |
| Anti-E clone 5G5 | 1:1000 (WB) | BEI resources [79] | Cat# NR-40318 |
| Anti-FLAG clone M2 | 1:1000 (WB) | Sigma | Cat# F1804 |
| Anti-rabbit IgG 680RD or 800CW | 1:20000 (WB) | Li-Cor Biosciences | Cat# 926-68071,<br>926-32213 |
| Anti-mouse IgG 680RD or 800CW | 1:20000 (WB) | Li-Cor Biosciences | Cat# 926-68070,<br>926-32210 |
| Anti-chicken IgY 800CW | 1:10000 (WB) | Li-Cor Biosciences | Cat# 926-32218 |
| Anti-rabbit AlexaFluor 488, 568, 647 | 1:1000 (IF) | Invitrogen | Cat# A11034,<br>A21069, A21246 |
| Anti-mouse AlexaFluor 568, 647 | 1:1000 (IF) | Invitrogen | Cat# A11031,<br>A21246 |
| Anti-chicken AlexaFluor 568, 647 | 1:1000 (IF) | Invitrogen | Cat# A11041,<br>A21449 |
| Streptavidin 680RD | 1:10000 (WB) | Li-Cor Biosciences | Cat# 68079 |

WB, Western blot; IF, immunofluorescence
\* The anti-E clone 5G5 was obtained from the Joel M. Dalrymple - Clarence J. Peters USAMRIID Antibody Collection through BEI Resources, NIAID, NIH: Monoclonal Anti-Langat Virus Envelope Glycoprotein (E), Clone 5G5 (produced *in vitro*), NR-40318.

**Table 3.**
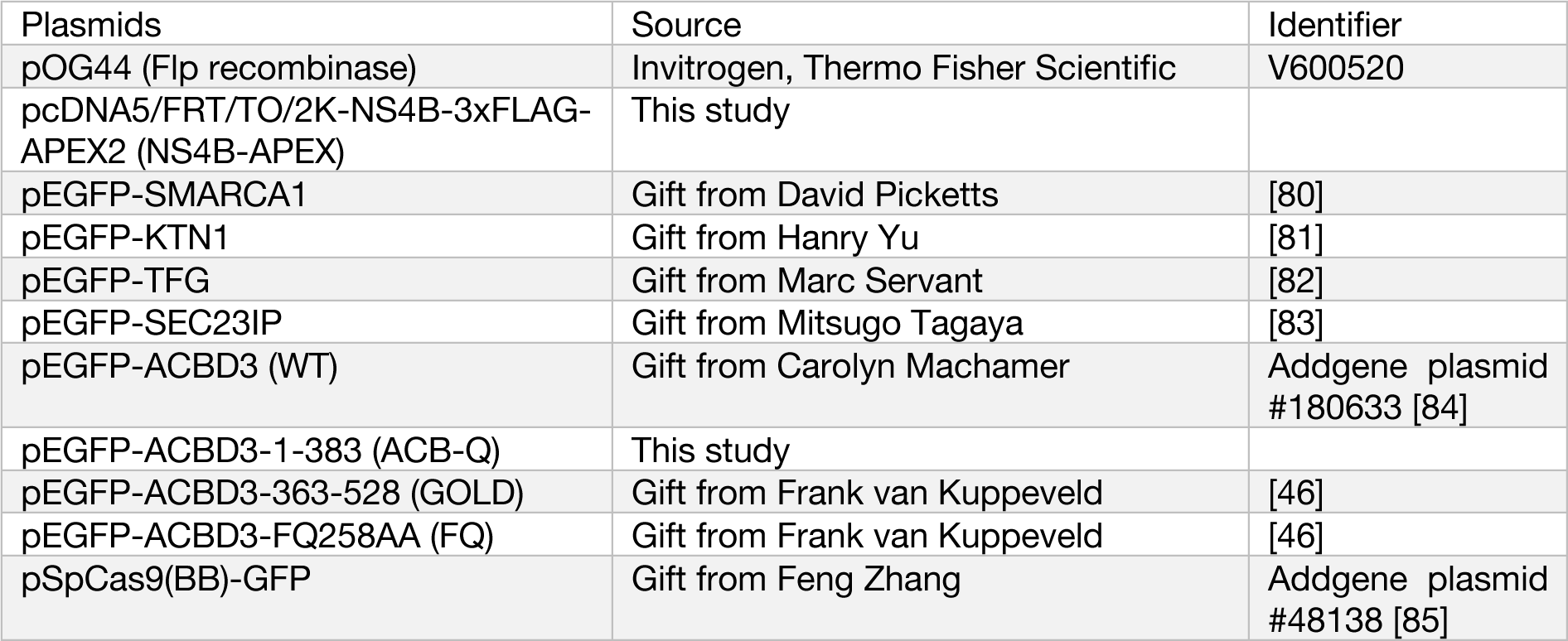
Plasmids used in the study.

### Generation of the NS4B-3xFLAG-APEX2 plasmid and the NS4B-APEX cell line

The coding sequence of APEX2 was PCR-amplified from APEX2 plasmid (a gift from Alice Ting, Addgene plasmid #72480, [86]) using specific primers containing restriction sites NotI (5’) and XhoI (3’). The amplified 5’ NotI-APEX2-XhoI 3’ fragments were digested and ligated into the vector pcDNA5/FRT/TO (Invitrogen, Thermo Fisher Scientific). Synthetic oligonucleotides encoding 3xFLAG were then ligated to the 5’ position of APEX2 sequence using NEBuilder HiFi DNA assembly kit (New England Biolabs) to form the final vector pcDNA5/FRT/TO/3xFLAG-APEX2.

Next, the coding sequence of 2K-NS4B was PCR-amplified from TBEV subgenomic DNA replicons [87] using specific primers containing restriction site KpnI (5’) and NotI (3’). Sequence of 2K was included to ensure the correct topology of NS4B at ER membrane. The amplified 5’ KpnI-2K-NS4B-NotI 3’ fragments were digested and ligated into the vector pcDNA5/FRT/TO/3xFLAG-APEX2 to form the plasmid NS4B-3xFLAG-APEX2 for the NS4B-APEX cell line.

Flp-In T-Rex HEK293 cells (Invitrogen, Thermo Fisher Scientific) were co-transfected with plasmids pOG44 (Invitrogen, Thermo Fisher Scientific) and pcDNA5/FRT/TO/2K-NS4B-3xFLAG-APEX2 in 9:1 (w/w) ratio using GeneJuice transfecting agent (Sigma-Merck) according to the manufacturer’s instructions. After 2 days of incubation, the cells were selected by 200 µg/ml hygromycin B (Invivogen) for 7 days. The cell line was maintained as above in medium containing 5 µg/ml of blasticidin (Invivogen) and 100 µg/ml of hygromycin B (Invivogen). Expression of NS4B-APEX protein was induced by incubating the cells with 1 ng/ml of doxycycline (Invitrogen, Thermo Fisher Scientific) for 16 hours.

### APEX biotinylation and pulldown

Confluent NS4B-APEX cells were incubated in 10 cm dishes with media containing 1 ng/ml doxycycline for 16 hours to induce expression of NS4B-APEX proteins. In case of LGTV infection, the cells were infected with LGTV for 2 hours prior to doxycycline induction. APEX-biotinylation was performed as previously published [33]. Briefly, the cells were treated with 50 µM biotin-phenol (Iris Biotech) for 30 min at 37°C. The cells were then incubated with 1 mM hydrogen peroxide for 1 min at room temperature to trigger biotinylation. The reaction was then quenched by washing the cells three times with ice-cold PBS buffer containing 5 mM Trolox (Sigma-Merck), 10 mM sodium ascorbate, and 10 mM sodium azide. For pulldown of biotinylated proteins, the cells were lysed with ice-cold RIPA buffer (25 mM HEPES pH 7.5, 150 mM NaCl, 1 mM EDTA, 1 mM EGTA, 1% NP-40, 1% sodium deoxycholate and 0.1% SDS) containing 5 mM Trolox, 10 mM sodium ascorbate, and 10 mM sodium azide, protease and phosphatase inhibitors (Roche). The lysates were centrifuged at 20,000 x g for 10 min at 4 °C. The cleared supernatants were incubated with 50 µl of neutravidin agarose bead slurry (Pierce, Thermo Fisher Scientific) at 4°C overnight. The beads were then washed once with each of the following buffers: RIPA buffer, 1 M KCl in H2O, 0.1 M Na2CO3 in H2O, 2 M urea in 10 mM Tris HCl, pH 7.4 and final 2% SDS in 150 mM NaCl with 25 mM HEPES pH 7.5. The washed beads were then incubated with 50 µl of 2x Laemmli buffer (without bromophenol blue and glycerol) containing 50 mM DTT and 5 mM biotin for 10 min at 75°C to elute the bound proteins. The samples were either analyzed with western blot or further processed for quantitative proteomic analysis.

### Quantitative proteomic analysis

#### Sample preparation

The APEX biotinylation pulldown samples were processed using modified filter-aided sample preparation (FASP) method [88,89]. In short, eluates were reduced with 100 mM DTT at 60°C for 30 min, transferred to Microcon-30kDa Centrifugal Filter Units (Merck), washed several times with 8 M urea and once with digestion buffer (DB, 50 mM TEAB, 0.5% sodium deoxycholate (SDC)) prior to alkylation with 10 mM methyl methanethiosulfonate in DB for 30 min in RT. Samples were digested with trypsin (0.3 µg Pierce MS grade Trypsin, Thermo Fisher Scientific) at 37°C overnight and an additional portion of trypsin (0.3 µg) was added, and the samples were incubated for another two hours. The samples were combined into one tandem mass tag (TMT) set and SDC was removed by acidification with 10% trifluoroacetic acid (TFA). The TMT-set was further purified using High Protein and Peptide Recovery Detergent Removal Spin Column and Pierce peptide desalting spin columns (both Thermo Fischer Scientific) according to the manufacturer’s instructions. Basic reverse phase peptide separation was performed using a Dionex Ultimate 3000 UPLC system (Thermo Fischer Scientific) and a reversed-phase XBridge BEH C18 column (3.5 μm, 3.0×150 mm, Waters Corporation) with a gradient from 3% to 100% acetonitrile in 10 mM ammonium formate at pH 10.00 over 27 min at a flow of 400 µL/min. The 10 fractions were dried and reconstituted in 3% acetonitrile, 0.1% TFA.

#### LC-MS analysis

The fractions were analysed on Orbitrap Fusion™ Tribrid™ mass spectrometer interfaced with nLC 1200 liquid chromatography system (both Thermo Fisher Scientific). Peptides were trapped on an Acclaim Pepmap 100 C18 trap column (100 μm x 2 cm, particle size 5 μm, Thermo Fischer Scientific) and separated on an in-house constructed analytical column (350 x 0.075 mm I.D.) packed with 3 μm Reprosil-Pur C18-AQ particles (Dr. Maisch, Germany) using a gradient from 3% to 80% acetonitrile in 0.2% formic acid over 90 min at a flow of 300 nL/min. Precursor ion mass spectra were acquired at 120,000 resolution, scan range 375-1375 and maximum injection time of 50 ms. MS2 analysis was performed in a data-dependent mode, where the most intense doubly or multiply charged precursors were isolated in the quadrupole with a 0.7 m/z isolation window and dynamic exclusion within 10 ppm for 60 s. The isolated precursors were fragmented by collision induced dissociation (CID) at 35% collision energy with the maximum injection time of 50 ms for 3 s (‘top speed’ setting) and detected in the ion trap, followed by multinotch (simultaneous) isolation of the top 10 MS2 fragment ions within the m/z range 400-1200, fragmentation (MS3) by higher-energy collision dissociation (HCD) at 65% collision energy and detection in the Orbitrap at 50 000 resolution m/z range 100-500 and maximum injection time 105 ms.

#### Database matching and quantitative analysis

The raw files were merged for identification and relative quantification using Proteome Discoverer version 2.4 (Thermo Fisher Scientific). The search was against SWISS-PROT *Homo sapiens* [90] and a custom database containing LGTV polyprotein and background proteins (using Mascot 2.5 (Matrix Science)) as a search engine with precursor mass tolerance of 5 ppm and fragment mass tolerance of 0.6 Da. Tryptic peptides were accepted with zero missed cleavage, variable modifications of methionine oxidation and fixed cysteine alkylation, TMT-label modifications of N-terminal and lysine were selected. Percolator was used for PSM validation with the strict FDR threshold of 1%. TMT reporter ions were identified with 3 milli mass units mass tolerance in the MS3 HCD spectra and no normalization was applied. Only the quantitative results for the unique peptide sequences with the minimum SPS match % of 55 and the average S/N above 10 were taken into account for the protein quantification. The quantified proteins were filtered at 1% FDR and grouped by sharing the same sequences to minimize redundancy.

### CRISPR-Cas9 knockdown and flow cytometry

The guide RNA (listed in table 4) was designed with the CRISPR design tool at Benchling (www.benchling.com) and cloned into vector pSpCas9(BB)-GFP (a gift from Feng Zhang, Addgene plasmid #48138) as described [85,91]. The cloned plasmids were transfected into HEK293T cells using GeneJuice transfecting agent (Merck Millipore). Forty-eight hours after transfection, the cells were re-seeded and transfected again for 48 hours. Following this the cells were then re-seeded into 12-well plates and infected with LGTV at MOI 1 for 2 hours at 37°C, 5% CO2. The inocula were then replaced with DMEM supplemented with 2% FBS. After 24 h.p.i., the cells were fixed in 4% formaldehyde for 20 min, and then washed three times with FACS buffer (20 mM EDTA, 2% FBS, 0.02% sodium azide in PBS) containing 0.5% Triton X-100. The cells were then incubated with anti-E antibodies in FACS buffer for 1 hour at RT. Then the cells were incubated with secondary antibodies AlexaFluor 647 for 1 hour at RT. The cells were then analyzed with a BD Accuri C6 flow cytometer (BD Biosciences).

**Table 4.**
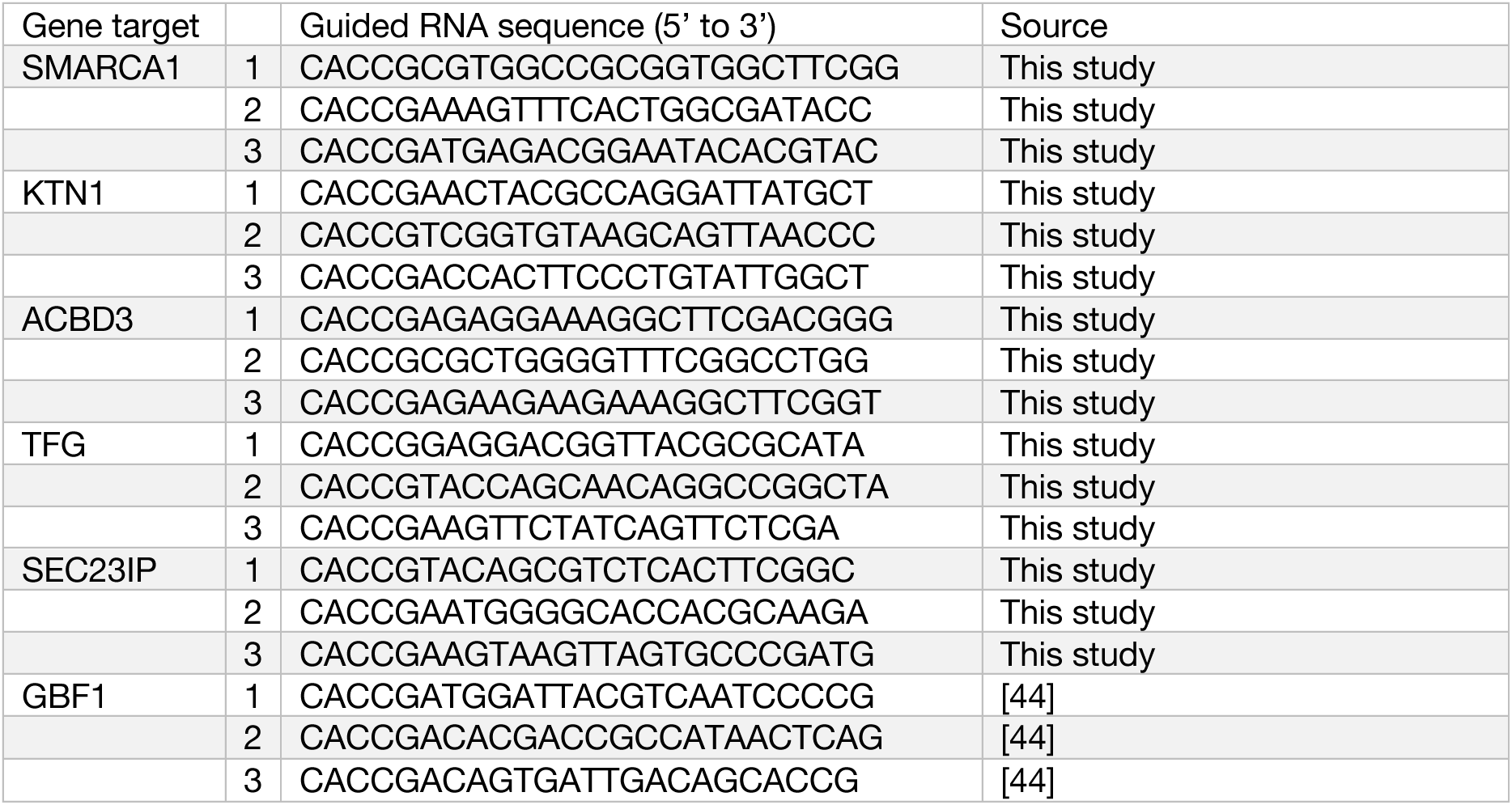
Guide RNA used in the study.

### Immunofluorescence staining

Cells on cover glasses were fixed with 4% formaldehyde for 20 min at RT and then rinsed with PBS. The fixed cells were then permeabilized with 0.1% Triton X-100 in PBS for 10 min at RT and then rinsed with PBS. The cells were then blocked with 2% BSA in PBS containing 0.05% Tween-20 (PBS-T) for 1 hour at RT. The cells were then stained with specific primary antibodies and secondary fluorescent antibodies and DAPI diluted in blocking buffer for 1 hour each. The fluorescent images were acquired by a Leica SP8 confocal microscope with a HC PL APO 63x/1.4 oil CS2 objective (Leica). For SIM, fluorescent images were acquired using a Zeiss Elyra 7 microscope with Lattice-SIM^2^ with a Plan-Apochromat 63x/1.4 oil objective (Zeiss).

### Quantification of confocal fluorescence microscopy images

Confocal fluorescence images were analyzed using ImageJ Fiji software [92]. NS3 regions in cells were determined by setting the fluorescent intensity threshold to 70. The NS3 regions of size > 10 µm^2^ were identified using the “Analyze Particles” function of ImageJ Fiji. The mean fluorescence intensities of NS3, E, or dsRNA within the identified NS3 regions were then measured. For the analysis of the fluorescence overlap, images were thresholded to generate a pixel area map of each fluorescent signal. Overlapping pixels in between different fluorescent signals were identified using “Image Calculator” function of ImageJ Fiji.

### Western blot

Cells were lysed in 2x Laemmli buffer (Bio-Rad) containing 100 mM DTT and heated for 10 min at 95°C. The cell lysate was then separated by electrophoresis in a 10% polyacrylamide gel and transferred to a PVDF membrane (Merck Millipore). The membrane was blocked with 5% milk in PBS-T. The blocked membrane was then incubated with primary antibodies for 1 hour at RT or 4°C overnight. The membrane was then incubated with secondary fluorescent antibodies (Li-Cor Biosciences) and detected with Odyssey Fc imager (Li-Cor Biosciences) and analyzed with Image Studio software (Li-Cor Biosciences).

### Virus infection quantification using plate imager

HEK293T cells and ACBD3 KO cells, were cultured in 96-well plates (Greiner Bio-one) and infected with indicated flaviviruses for 2 hours at 37°C, 5% CO2. MOI of the viruses used are described in the figure legends. The inocula were then replaced with DMEM supplemented with 2% FBS. After 24 h.p.i., the cells were fixed in 4% formaldehyde for 20 min, and then permeabilized with 0.5% Triton X-100 in PBS. The cells were then incubated with either anti-E antibodies or anti-dsRNA antibodies (specified in the figure legends) for 1 hour at RT. Then the cells were incubated with secondary antibodies AlexaFluor 647 and DAPI for 1 hour at RT. The plates were then analyzed with a Cytation 5 plate imager with Gen5 software (Biotek) to quantify the number of either E protein or dsRNA positive cells.

### PI4KB inhibition assay

A549 cells were cultured in 96-well plates (Greiner Bio-one) and infected with indicated flaviviruses at MOI 0.1 (0.5 for DENV2) for 2 hours at 37°C, 5% CO2. For PV, infection with PV at MOI 10 for 1 hour at 37°C, 5% CO2. The inocula were then replaced with DMEM supplemented with 2% FBS. PI4KB inhibitor T-00127-HEV1 (Calbiochem # 538001, CAS

900874-91-1) or DMSO control was added to the media. After 24 h.p.i. (6 h.p.i. for PV), the cells were fixed in 4% formaldehyde for 20 min, and then permeabilized with 0.5% Triton X-100 in PBS. The cells were then incubated with anti-dsRNA antibodies for 1 hour at RT. Then the cells were incubated with secondary antibodies AlexaFluor 647 and DAPI for 1 hour at RT. The plates were then analyzed with a Cytation 5 plate imager with Gen5 software (Biotek) to quantify the number of E protein positive cells.

### Rescue assay with ACBD3 mutants

HEK293T cells and ACBD3 KO cells were transiently transfected with either GFP or the indicated GFP-ACBD3 mutants for 24 hours. The transfected cells were then infected with TBEV at MOI 0.1 for 2 hours at 37°C, 5% CO2. The inocula were then replaced with DMEM supplemented with 2% FBS. After 24 h.p.i., the cells were fixed in 4% formaldehyde for 20 min, and then permeabilized with 0.5% Triton X-100 in PBS. The cells were then incubated with anti-E antibodies for 1 hour at RT. Then the cells were incubated with secondary antibodies AlexaFluor 647 and DAPI for 1 hour at RT. The plates were then analyzed with a Cytation 5 plate imager with Gen5 software (Biotek) to quantify the number of GFP positive and E protein positive cells.

### Quantitative reverse-transcription PCR (RT-qPCR)

Total RNA of LGTV-infected cells was extracted using NucleoSpin RNA Plus kit (Macherey-Nagel) and eluted in 60 µl RNase-free H2O according to the manufacturer’s instructions. Ten μl of total RNA extract was used for cDNA synthesis using High Capacity cDNA Reverse Transcription kit (Thermo Fisher Scientific) according to manufacturer’s instructions. LGTV RNA was quantified using qPCRBIO probe mix Hi-ROX (PCR Biosystems) and primers recognizing NS3, forward primer 5′-AACGGAGCCATAGCCAGTGA-3′, reverse primer 5′-AACCCGTCCCGCCACTC-3′ and probe FAM-AGAGACAGATCCCTGATGG-BHQ. Actin was used as a housekeeping gene to normalize the amount of viral RNA by using the qPCRBIO SyGreen mix HI-ROX (PCR Biosystems) together with QuantiTect primer assay (QT01680476, Qiagen). All Quantitative PCR (qPCR) experiments were performed on the StepOnePlus fast-real-time PCR system (Applied Biosystems).

### Virus-like particle secretion assay

HEK293T cells and ACBD3 KO cells were grown in T75 flasks and transfected for 24 hours with 3 µg of plasmids encoding for viral proteins C, prM and E-3xFLAG each [9], or 9 µg of GFP plasmid as control, using GeneJuice transfecting agent (Sigma-Merck) according to the manufacturer’s instructions. The culture was then harvested as cells and supernatant separately. First, cells were detached using cold PBS and lysed in 350 µl lysis buffer (50 mM Tris pH 8.0, 150 mM NaCl, 1% Triton X-100) containing protease inhibitor (Roche) for at least 20 min on ice. Cell lysate was sonicated for 7 seconds (Hielscher UP200St) and centrifuged at 20,000 x g for 10 min at 4 °C. Cleared lysate was transferred to a new tube, mixed with 2x Laemmli buffer (Bio-Rad) containing 100 mM DTT and boiled for 5 min at 95°C for western blot. Second, the culture supernatant harvested was clarified by centrifugation at 500 x g for 5 min at 4°C. Pre-cleared supernatant was ultracentrifuged at 100,000 x g for 1.5 hours at 4°C. The supernatant was then removed. The pellet was resuspended in 50 µl 2x Laemmli buffer containing 100 mM DTT overnight at 4°C. The samples were boiled for 5 min at 95 °C and analyzed with western blot as above. Quantification was performed by normalizing the protein in the VLPs to the protein in the lysate.

### Statistics

For the data of NS4B-APEX proteomics, statistical significance was calculated using Excel (Microsoft). Statistical analyses of the other experiments were performed using Prism 10 (Graphpad). Unpaired t-test was used to analyze the data of plate imager-based assays, focus forming assays, RT-qPCR, and VLP secretion assay. Mann-Whitney test was used to analyze the data of confocal images. One-way ANOVA with Dunnett multiple test was used to analyze the data of CRISPR-Cas9 KD assay and rescue assay with ACBD3 mutants. The specific tests used and the p-values are indicated in the figure legends.

## Data availability

Proteomics data (dataset identifier PXD051173) are deposited at the ProtomeXchange Consortium via PRIDE partner repository [93].

## Acknowledgements

We thank Frank van Kuppeveld (Utrecht University) for the ACBD3 KO cells and the ACBD3 mutant plasmids, David Picketts (Ottawa University) for the SMARCA1 plasmid, Hanry Yu (National University of Singapore) for the KTN1 plasmid, Carolyn Machamer (Johns Hopkins University) for the ACBD3 plasmid, Mitsugo Tagaya (Tokyo University of Pharmacy and Life Sciences) for the SEC23IP plasmid, and Marc Servant (Montreal University) for the TFG plasmid. We acknowledge the Biochemical Imaging Center (BICU) at Umeå University and the National Microscopy Infrastructure, NMI (VR-RFI 2016-00968) for providing assistance in microscopy. Proteomic analysis was performed at the Proteomics Core Facility, Sahlgrenska academy, Gothenburg University, with financial support from SciLifeLab and BioMS. We thank Annika Thorsell (Proteomics Core Facility, Gothenburg University) for the proteomics data and analysis. The work was supported by grants from the Swedish Research Council (2021-05117 and 2018-05851) to RL and (2018-05851 and 2020-06224) to AKÖ. Open access funding provided by Umeå University.

## Author contributions

### Conceptualization

Wai-Lok Yau, Richard Lundmark

### Data curation

Wai-Lok Yau, Marie Berit Akpiroro Peters

### Formal analysis

Wai-Lok Yau, Marie Berit Akpiroro Peters, Marie Sorin, Sebastian Rönfeldt, Richard Lundmark

### Funding acquisition

Lars-Anders Carlson, Anna K Överby, Richard Lundmark

### Investigation

Wai-Lok Yau, Marie Berit Akpiroro Peters, Marie Sorin, Sebastian Rönfeldt, Richard Lundmark

### Project administration

Richard Lundmark, Anna K Överby

### Supervision

Lars-Anders Carlson, Anna K Överby, Richard Lundmark

### Visualization

Wai-Lok Yau, Richard Lundmark

### Writing – original draft

Wai-Lok Yau, Lauri I.A. Pulkkinen, Richard Lundmark

### Writing – review & editing

Wai-Lok Yau, Marie Berit Akpiroro Peters, Marie Sorin, Sebastian Rönfeldt, Lauri I.A. Pulkkinen, Lars-Anders Carlson, Anna K Överby, Richard Lundmark

## Supporting information

**S1 Table. Mass spectrometry result and gene ontology analysis.**

STRING functional enrichment analysis of the NS4B-APEX proteomes was performed using STRING version 12.0 (https://string-db.org/) [94]. Small geneset-based analysis of protein input with FC value of hits of NS4B-APEX infected with LGTV over mock was used and compared with the whole human proteome and confidence of interaction score 0.7 was set. Venn diagram of the hits in S1 Table C was performed using BioVenn web application (https://www.biovenn.nl/index.php) [95]. ShinyGO enrichment analysis was performed using ShinyGO version 0.77 (http://bioinformatics.sdstate.edu/go/) [96] with the parameters p-value of FDR cutoff at 0.05, 10 pathways to show, pathway size from 2 to 100, remove redundancy and abbreviate pathways.

**S1 Fig.**
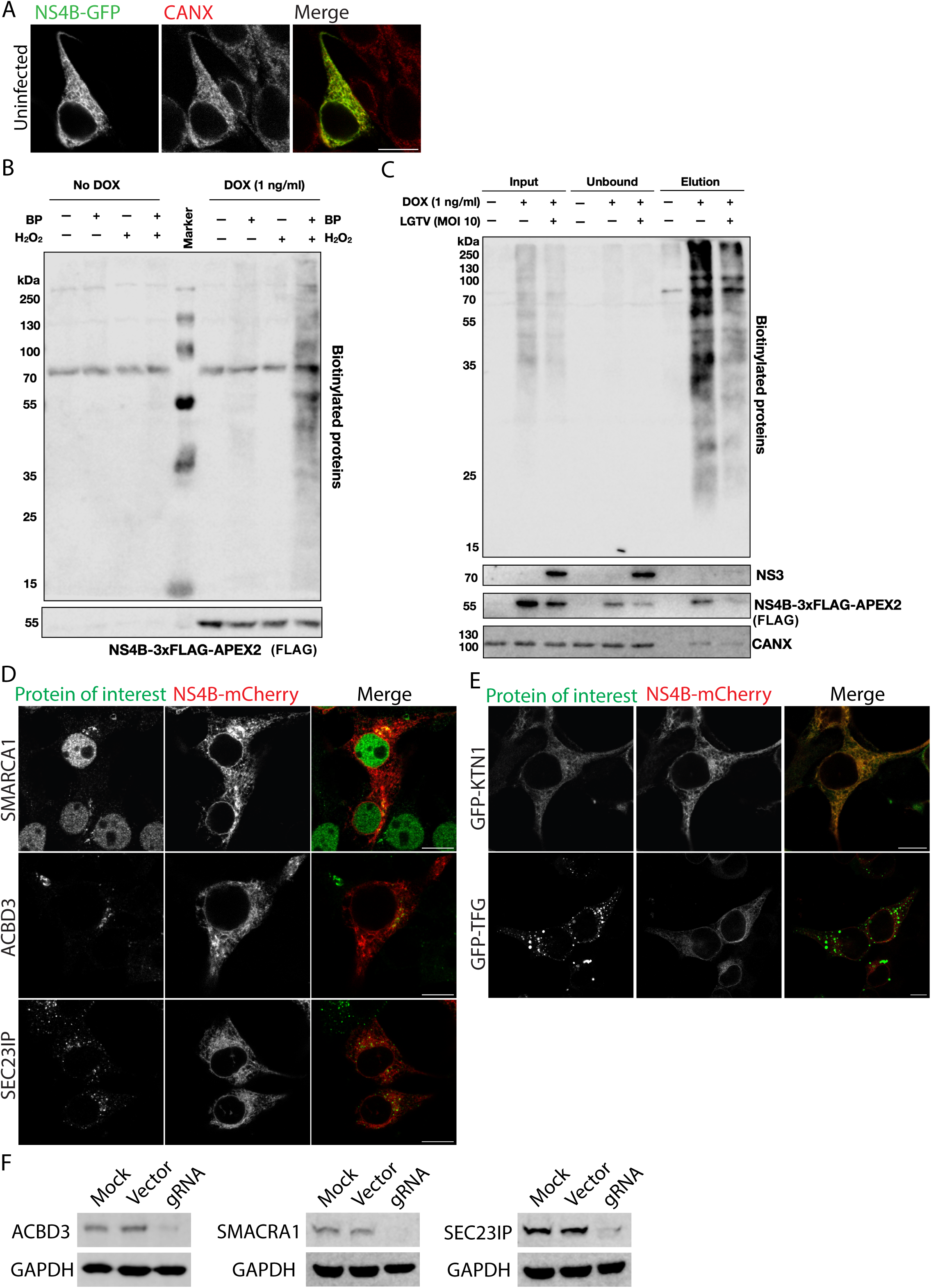
(related to Fig 1). Validation of NS4B-APEX proximal protein analysis. (A) Confocal fluorescence micrographs of uninfected HEK293T cells transiently over-expressing NS4B-GFP and stained with anti-CANX antibodies. Scale bar, 10 µm. (B) Western blot analysis of NS4B-APEX cells treated for APEX biotinylation under specified condition. The cell lysate was probed with streptavidin 680RD for biotinylated proteins and anti-FLAG M2 antibodies for NS4B-3xFLAG-APEX2. BP, substrate biotin phenol; H2O2, catalyst hydrogen peroxide; DOX, doxycycline. (C) Pulldown analysis of LGTV-infected (MOI 10, 16 h.p.i.) and doxycycline-induced NS4B-APEX cells treated for APEX biotinylation. The cell lysate was mixed with neutravidin beads to pulldown biotinylated proteins. Cell lysate, unbound samples and elute were analyzed by western blot with streptavidin 680RD, anti-NS3, anti-CANX, anti-FLAG M2 antibodies. (D) Confocal fluorescence micrographs of HEK293T cells transiently expressing NS4B-mCherry and stained with either anti-SMARCA1, anti-ACBD3 or anti-SEC23IP antibodies. Scale bar, 10 µm. (E) Confocal fluorescence micrograph of HEK293T cells transiently co-expressing NS4B-mCherry with either GFP-KTN1 or GFP-TFG. Scale bar, 10 µm. (F) Analysis of CRISPR-Cas9 KD efficiency of ACBD3, SMARCA1 and SEC23IP by western blot with specific antibodies. GAPDH was used as loading control of the samples.

**S2 Fig.**
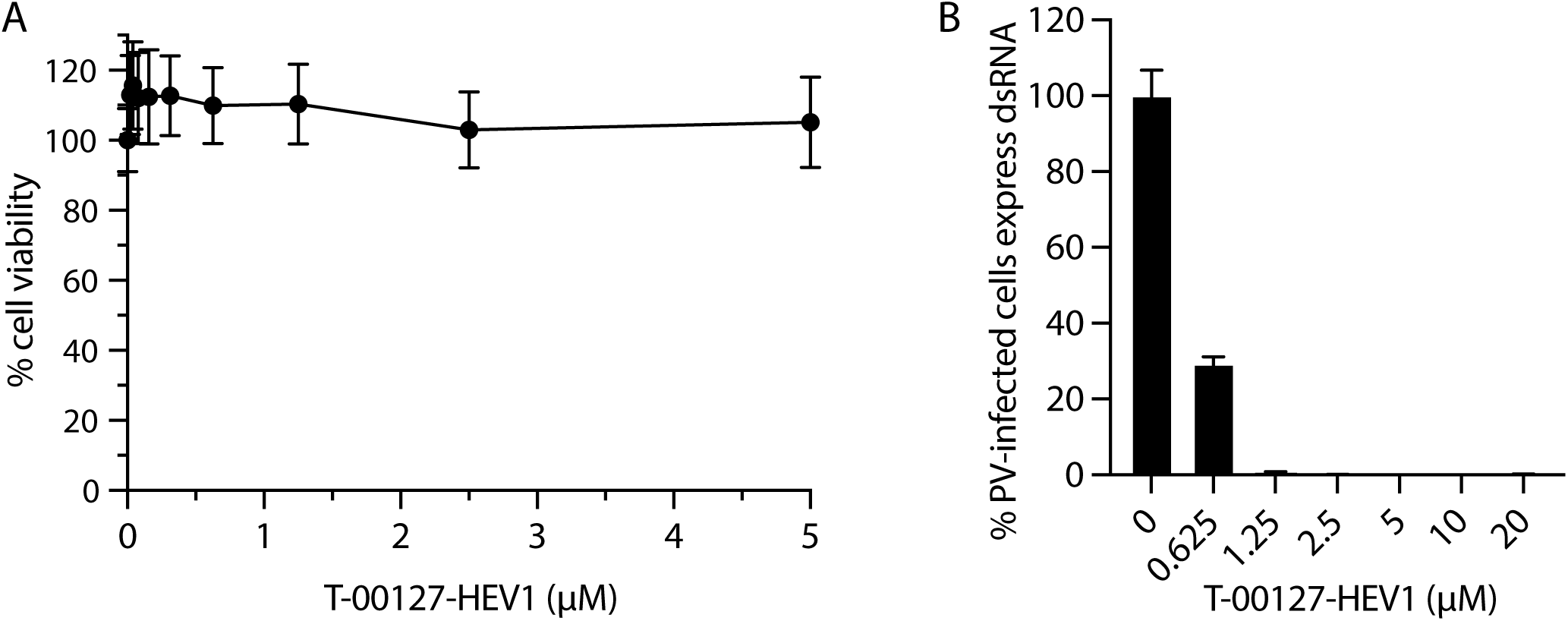
(related to Fig 2). PI4KB inhibitor T-00127-HEV1 inhibits poliovirus infection. (A) Viability test of A549 cells treated with PI4KB inhibitor T-00127-HEV1. A549 cells were treated with T-00127-HEV1 for 24 hours at 37°C with 5% CO2. Cell viability of the cells was determined with Celltiter Glo luminescent cell viability assay (Promega) and Synergy HT plate reader (Biotek). Mean ± SD of 4 biological replicates from a representative experiment. (B) Quantification of the percentage of A549 cells positive for dsRNA at 6 h.p.i. with poliovirus (MOI 10), in the absence and presence of T-00127-HEV1 (0 – 20 µM). Data were normalized to DMSO-treated A549 control cells. Mean ± SD of 4 biological replicates from a representative experiment.

**S3 Fig.**
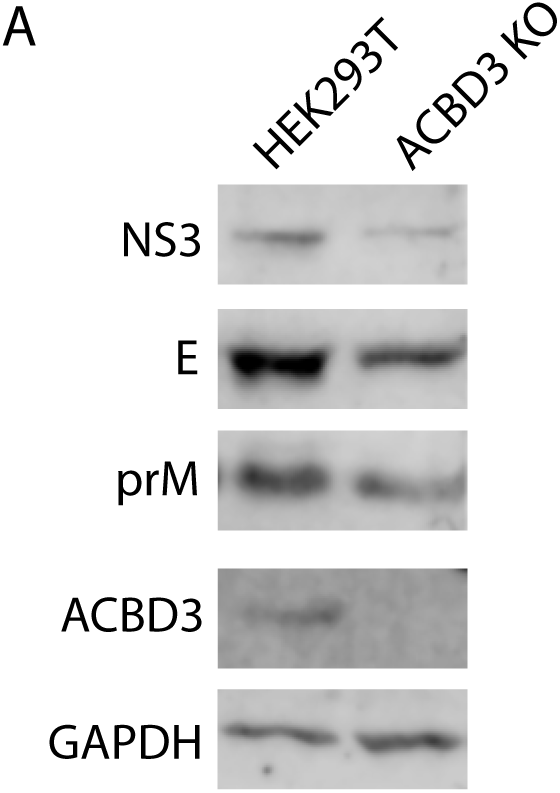
(related to Fig 3). ACBD3 KO inhibits LGTV protein expression. (A) Expression of viral proteins of LGTV-infected (MOI 1, 24 h.p.i.) HEK293T cells and ACBD3 KO cells. The cell lysate was analyzed by western blot with specific antibodies to NS3, E, prM, ACBD3 and GAPDH.

**S5 Fig.**
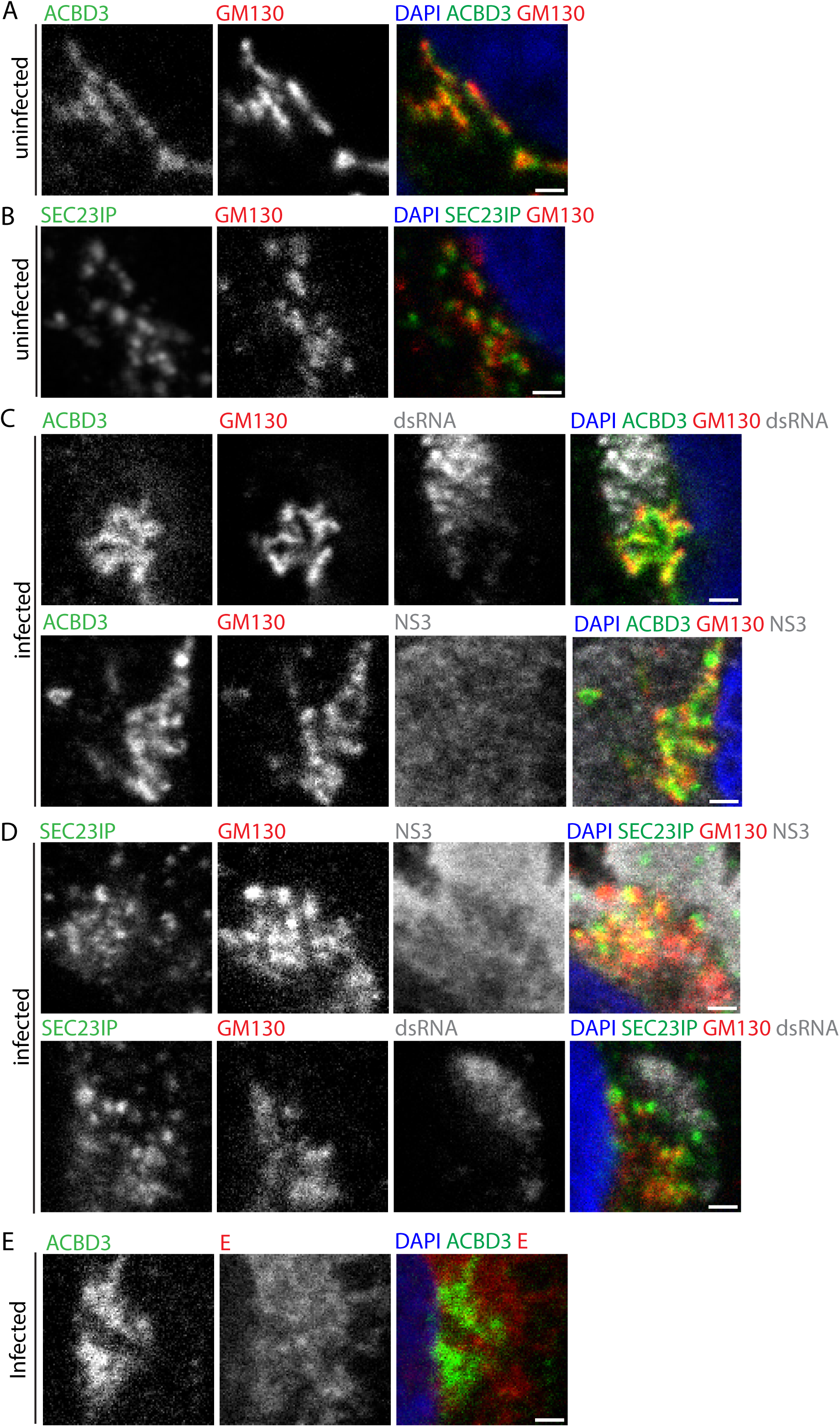
(related to Fig 5). ACBD3 and SEC23IP localize at the ERES-Golgi contact. (A) Confocal fluorescence micrographs of uninfected HEK293T cells stained with anti-ACBD3 antibodies and anti-GM130 antibodies. Scale bar, 1 µm. (B) Confocal fluorescence micrograph of uninfected HEK293T cells stained with anti-SEC23IP antibodies and anti-GM130 antibodies. Scale bar, 1 µm. (C) Confocal fluorescence micrographs of LGTV-infected (MOI 1, 16 h.p.i.) HEK293T cells and stained with specific antibodies to ACBD3, GM130, dsRNA and NS3 as indicated. Scale bar, 1 µm. (D) Confocal fluorescence micrographs of LGTV-infected (MOI 1, 16 h.p.i.) HEK293T cells stained with specific antibodies to SEC23IP, GM130, NS3 and dsRNA as indicated. Scale bar, 1 µm. (E) Confocal fluorescence micrographs of LGTV-infected (MOI 1, 16 h.p.i.) HEK293T cells stained with anti-ACBD3 antibodies and anti-E antibodies. Scale bar, 1 µm.

**S6 Fig.**
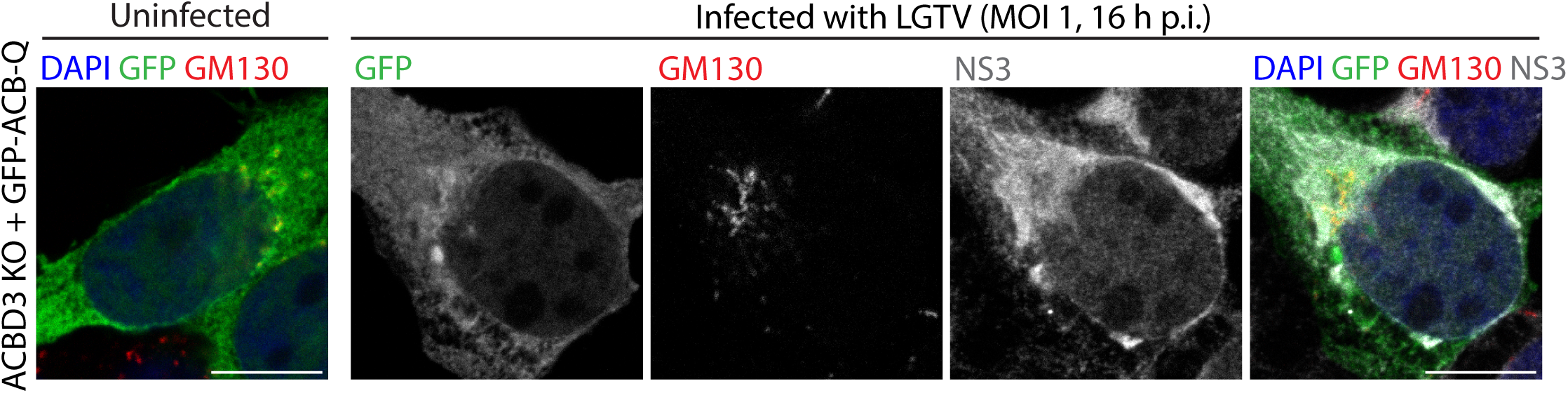
(related to Fig 6). ACBD3 ACB-Q domain localizes in the cytoplasm. Confocal fluorescence micrographs of ACBD3 KO cells transiently expressing GFP-ACB-Q at 16 h.p.i. with LGTV (MOI 1) and stained with anti-GM130 antibodies and anti-NS3 antibodies. Scale bar, 10 µm.

**S1 movies (related to Fig 5). 3D representations of the volume SIM imaging data of ACBD3 and SEC23IP in uninfected cells.**

HEK293T cells transiently expressing GFP-ACBD3 were stained anti-SEC23IP antibodies. (A) Movie of the Z-stack of the SIM images; (B) Movie of the inset; (C) Movie of the volume SIM imaging data of the inset. Green, GFP-ACBD3; Red, SEC23IP.

**S2 movies (related to Fig 5). 3D representations of the volume SIM imaging data of ACBD3 and SEC23IP in infected cells.**

HEK293T cells transiently expressing GFP-ACBD3 at 16 h.p.i. with LGTV (MOI 1) were stained anti-SEC23IP antibodies and anti-E antibodies. (A) Movie of the Z-stack of the SIM images; (B) Movie of the inset; (C) Movie of the volume SIM imaging data of the inset. Green, GFP-ACBD3; Red, SEC23IP; Gray, E.

